# Real-Time Point Process Filter for Multidimensional Decoding Problems Using Mixture Models

**DOI:** 10.1101/505289

**Authors:** Ali Yousefi, Mohammad Reza Rezaei, Kensuke Arai, Loren M. Frank, Uri T. Eden

**Affiliations:** Department of Mathematics and Statistics, Boston University, Boston, MA 02215; Department of Electrical and Computer Engineering, Isfahan University of Technology, Isfahan 84156-83111, Iran; Department of Physiology, University of California, San Francisco, San Francisco, CA 94158

**Keywords:** Point-process filter, Real-time filter, Marked point-process filter, State-space modeling, Mixture model, Gaussian mixture model, Mixture merging algorithm, Mixture dropping algorithm

## Abstract

There is an increasing demand for a computationally efficient and accurate point process filter solution for real-time decoding of population spiking activity in multidimensional spaces. Real-time tools for neural data analysis, specifically real-time neural decoding solutions open doors for developing experiments in a closed-loop setting and more versatile brain-machine interfaces. Over the past decade, the point process filter has been successfully applied in the decoding of behavioral and biological signals using spiking activity of an ensemble of cells; however, the filter solution is computationally expensive in multi-dimensional filtering problems. Here, we propose an approximate filter solution for a general point-process filter problem when the conditional intensity of a cell’s spiking activity is characterized using a Mixture of Gaussians. We propose the filter solution for a broader class of point process observation called marked point-process, which encompasses both clustered – mainly, called sorted – and clusterless – generally called unsorted or raw– spiking activity. We assume that the posterior distribution on each filtering time-step can be approximated using a Gaussian Mixture Model and propose a computationally efficient algorithm to estimate the optimal number of mixture components and their corresponding weights, mean, and covariance estimates. This algorithm provides a real-time solution for multi-dimensional point-process filter problem and attains accuracy comparable to the exact solution. Our solution takes advantage of mixture dropping and merging algorithms, which collectively control the growth of mixture components on each filtering time-step. We apply this methodology in decoding a rat’s position in both 1-D and 2-D spaces using clusterless spiking data of an ensemble of rat hippocampus place cells. The approximate solution in 1-D and 2-D decoding is more than 20 and 4,000 times faster than the exact solution, while their accuracy in decoding a rat position only drops by less than 9% and 4% in RMSE and 95% HPD coverage performance metrics. Though the marked-point filter solution is better suited for real-time decoding problems, we discuss how the filter solution can be applied to sorted spike data to better reflect the proposed methodology versatility.

## 1. Introduction

The point-process modeling framework is widely used in the analysis of neural spike trains [1-5], and particularly in conjunction with the filtering framework [6, 7], can be used to link the spike train data to low-dimensional dynamical external covariates like movement or sensory inputs [2, 4, 5]. As examples, statistical filtering employing point process noise models has been used in estimation of a rat’s position given the spiking activity of its hippocampal place cells [2, 3], and also in decoding of arm-movements [8]. However, when the dimension of the dynamical covariates increases, computational cost becomes expensive [1, 9]. We previously developed a computationally efficient point-process filter solution for high-dimensional decoding problems, which reduces the computational cost of the filter implementation [10]. In the development of this algorithm, we used a non-parametric Gaussian kernel to estimate the conditional intensity functions of each cell from the data [11]. However, this estimation is still computationally expensive for real-time applications, which limits the application of the proposed filter solution in real-time experimental designs. In a more recent work, we have demonstrated the feasibility of building an adaptive parametric conditional intensity function (CIF) using Mixture of Gaussians (MoGs), similar to a Gaussian Mixture Model (GMM) for a distribution. MoG is a powerful and flexible model for representing multi-modal and complex functions and distributions and MoGs follows the same functional form as GMM, except its mixing weights are not normalized to one. Using this class of CIFs, we can develop new solutions which greatly reduce the computational cost of the filter solution. Here we demonstrate such a solution and present an accurate, computationally inexpensive point-process filter solution when neural activity of a cell is characterized by MoGs.

The point process filter is comprised of two models: the state transition model and an observation model [11, 12]. The state transition model characterizes how the external or behavioral covariate(s) – called the state variable – change over time, and the observation model defines the likelihood of observing a spike event as a function of the previous spiking activity and the state variable [12]. The state transition in the case of navigation through space can be well characterized by a low-dimensional random walk model [13-15]. Under this modeling assumption, for a posterior distribution in the GMM class, the one-step prediction [16] is again of GMM class, which has an analytical solution. The posterior distribution, e.g. the filter solution, is proportional to the product of one-step prediction and the likelihood of observed spiking data. Given a MoGs model for the cells’ conditional intensities (CIFs), the posterior can be thought of as a multiplication of two mixtures of Gaussian functions – one from the observation likelihood and the other one from one-step prediction. Multiplication of a MoGs and a GMM results in a new MoGs, to which a GMM is proportional to [17]; thus, the modeling challenge in this filter problem switches from calculating one-step filter and approximating the likelihood function [10] to optimally controlling the growth of the number of mixture components over each processing time-step.

There are two main approaches that are developed to manage the growth of number of mixture components. The first approach relies on functional approximation, where the filter solution at each time-step is approximated using a fixed number of mixture components [18-20]. The second approach relies on minimizing some forms of distance measure between two distributions [21-24]. In this approach, two GMMs, one which is completely known and the other one with a lower number of mixture components is built to approximate the first one. In practice, the first approach has a larger computational cost and induces bias in the filter solution as the pre-defined number of the mixture components or their fixed parameters can be optimal for a limited number of processing time-steps. The second approach has the capacity to change the number of mixtures and their parameters per each time-step given the observation process and its likelihood function [25]. However, there are a couple of modeling challenges in defining a proper distance measure, identifying an optimal number of mixtures, or finding an optimal stopping criterion. In this research, we mainly focus on the second approach and propose a revised distance measure and stopping criterion, which will address issues of the previously developed methodologies [5, 11].

In our solution, we use a symmetric Kullback–Leibler (KL) [26-28] distance and develop procedures for merging and dropping mixture components [26]. In the dropping procedure, we identify which mixture component(s) can be dropped from the mixture pool; whilst, in the merging procedure, we propose a new procedure to sequentially merge mixture components [29-31]. For both procedures, we define stopping criteria to determine when a further merging or dropping of mixture components is not desirable. We use both merging and dropping procedures in our filter solution and demonstrate this new filter solution application in both 1-D and 2-D decoding. We compare performance result of this approximate filter solution with the exact filter solution and show the computational time for both methods. In the appendix section, we also provide a comparison study between the classical KL measure and the symmetric one utilized in this research. This comparison highlights the importance of a proper distance measure in controlling the mixture growth. The drop-merge method proposed here can be applied to other non-linear and multi-modal filter problems when the computational cost of the exact solution becomes prohibitive.

We develop the filter solution for the marked point-process data. The marked point-process is a broader class of point-process, in which, each event in time has an associated mark. Unsorted or raw spiking data – broadly, called clusterless data – fall to the category of marked point-process, where features associated with each spike event define its mark. A sorted spike is a specific category of the marked point-process, where the identity of each spike is its mark information. The filter solution proposed here encompasses both clustered and clusterless spike data. We start by Methods section where we propose our filter solution. We then demonstrate the solution application in 1-D and 2-D decoding problem, where we provide an inclusive analysis of both performance and computational complexity of the proposed solution. In particular, we compare the solution with the numerical solution which is generally called the exact solution. We also provide a detailed analysis of the proposed solution in the appendix section. In the discussion section, we elaborate on the potential and application of our proposed solution along the future direction of this research.

## 2. Methods

### 2.1 Problem Definition

For the marked point-process observation, the instantaneous probability of observing a spike at time *t*with a spike waveform mark in ℳ – ℳ ∈ ℜ^*K*^, where *K* represents the dimension, is defined by

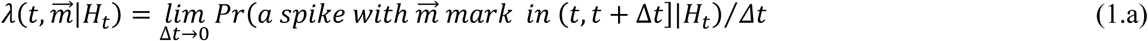

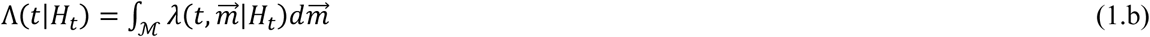

where 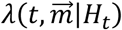 is called the joint mark intensity function, which defines the instantaneous probability of a spike at time *t* with a mark 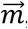, and Λ(*t*|*H*_*t*_) defines the intensity function of the “ground process” [11]. *H*_*t*_represents the full history of spiking from all recorded neurons up to time *t*[32, 33]. We assume neural spiking depends on a covariate vector *X*_*t*_; therefore, we construct a joint mark intensity model of the form 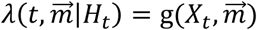. We further assume that the joint mark intensity function can be expressed as a MoGs over both *X*_*t*_and mark spaces

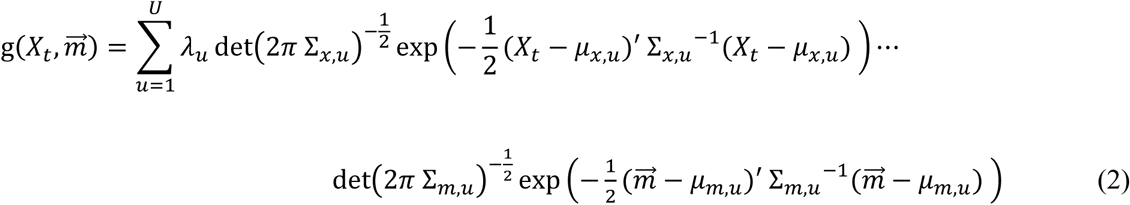

where, *λ*_*u*_ > 0 defines the rate of *u*^*th*^ mixture. Note that the sum of *λ*_*u*_ is not normalized. (*μ*_*x,u*_, ∑_*x,u*_) is the mean and covariance matrix of the *u*^*th*^ mixture model over *X*_*t*_space, and (*μ*_*m,u*_, ∑_*m,u*_) define the mean and covariance matrix of the corresponding mixture over mark space. Under this assumption, the intensity function of the ground process [9, 34] is

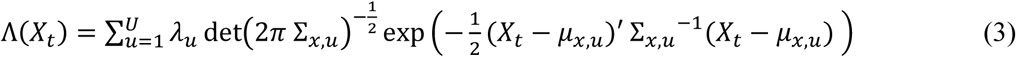

In the marked point-process framework, the likelihood of being at a coordinate *X*_*k*_ when observing *N*_*k*_, – assumed to be either 0 or 1, spikes in the interval Δ_*k*_= (*t*_*k*-1_, *t*_*k*_] with marks 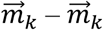 is observed when *N*_*k*_ is 1 – is defined by

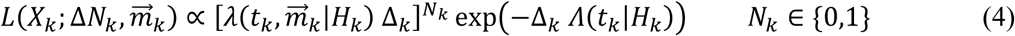

where, *X*_*k*_ is a discrete-time representation of *X*_*t*_ for the Δ_*k*_ time interval. Note that when there is more than one spike event in the Δ_*k*_ time interval, we can partition this interval to smaller time intervals and assume each of these multiple events happens in one of the shorter time intervals [11]. In equation (4), the joint mark conditional intensity and ground conditional intensity definition characterize neural activity of an ensemble of cells. For example, both models can represent multi-unit spike mark events recorded using a tetrode [35]. In Appendix A, we define the likelihood function for multiple ensembles of cells activity sampled from multiple independent groups of electrodes. In the rest of this paper, we focus on the likelihood function defined in equation (4). Extension of the filter solution and drop-merge algorithms to models of activity of multiple cell ensembles is straight forward.

Using 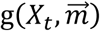 and Λ(*X*_*t*_) definition, the likelihood function is defined by

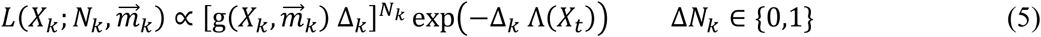

We assume that the time *X*_*k*_ evolution is a Markovian process [36]. We also assume that the time evolution of *X*_*k*_ can be described by a linear state equation

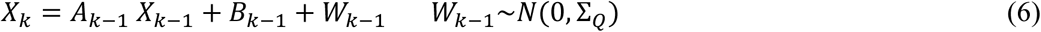

where *W*_*k*_ is a multivariate normal with zero mean, and *A*_*k*-1_ and *B*_*k*-1_ are the state and input matrices defining the state evolution over time. Under this assumption, the one-step density of state *X*_*k*_, is defined by

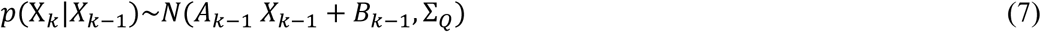

Given the observation process and the state evolution equation, the exact posterior distribution of the state at time index *k* is defined by

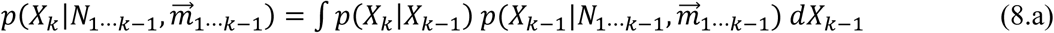

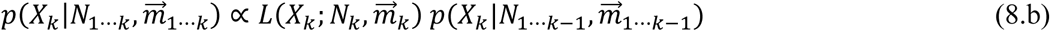

We assume 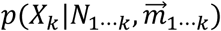 can be approximated by a GMM defined by

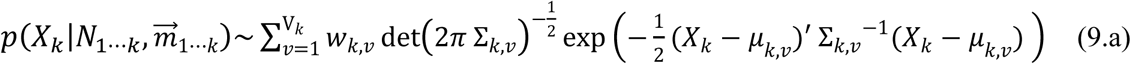

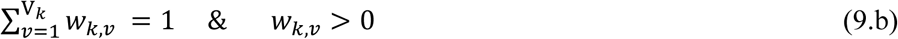

where, V_*k*_ is the number of mixture components at time index *k* and *w*_*k,v*_ is the weight of the *v*^*th*^ mixture component with corresponding mean and covariance matrices – (*μ*_*k,v*_, ∑_*k,v*_).

Using those definitions, we now develop a solution that estimates a parsimonious number of mixture components V_*k*_, and corresponding (*μ*_*k,v*_, ∑_*k,v*_) per each filtering time-step. Under the assumption that Λ(·) is constant over the state domain, there is also a closed-form solution for the one-step prediction (equation 8.a) and filter update (equation 8.b). Appendix B describes the closed form solution for the general case where Λ(*t*|*H*_*t*_) is not constant in detail. However, the number of mixture components is multiplied by a scaler factor corresponding to the number of mixture components in the likelihood function – equation (8.b) – per each filtering time-step and this leads to an exponential growth in the number of mixture components over time, as does the computational cost of the filter solution. Fortunately, many of these mixture components either have an infinitesimal weight or share a similar mean and covariance matrices, allowing us to optimally drop or merge mixture components on each filtering step. Here, we demonstrate how mixtures merging and dropping algorithms, which are later used in 1-D and 2-D decoding of rat position in a maze using neural spiking activity recorded from multiple tetrodes.

### 2.2 Dropping and Merging Procedure for a GMM

Let’s assume *P* is a GMM defined by

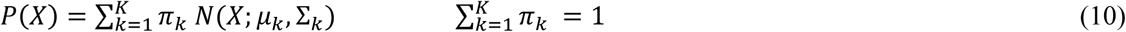

where *K* is the number of mixture components and *π*_*k*_ is the mixing weight for the *k*^*th*^ mixture component. For this GMM, (*μ*_*k*_, ∑_*k*_) define the mean and covariance of the *k*^*th*^ mixture component. We want to merge or drop *P* components, while the new mixture model *Q* properly represents *P*. We use the following divergence measure to assess the similarity between *P* and *Q* distributions, which is defined by

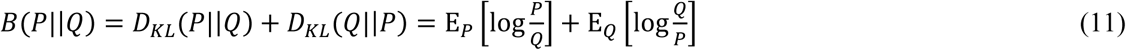

where, *D*_*KL*_(*P*||*Q*) is KL divergence from *P* to *Q*, and *D*_*KL*_(*Q*||*P*) is KL divergence from *Q* to *P*. *B*(*P*||*Q*) is nonnegative and symmetric, and it is zero when *P* and *Q* are the same [28]. Thus, the objective is to minimize *B*(·) while the components of *P* are merged or dropped.

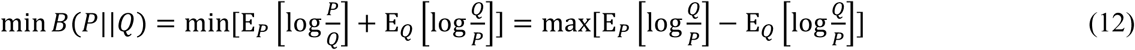

The idea behind constructing the *B*(*P*||*Q*) divergence measure is that the first term – *D*_*KL*_(*P*||*Q*) – favors similarity between *Q* and *P* over the spaces covered by *P* components, and the second term – *D*_*KL*_(*Q*||*P*) – punishes the same similarity over the spaces covered by *Q* components. While *D*_*KL*_(*P*||*Q*) is less sensitive to merging components of *P* in constructing *Q, D*_*KL*_(*Q*||*P*) favors keeping components of *P*, specifically those whose merging causes the space covered by *Q* to grow. We can also change contribution of *D*_*KL*_(*P*||*Q*) and *D*_*KL*_(*Q*||*P*) in *B*(*P*||*Q*), which is defined by

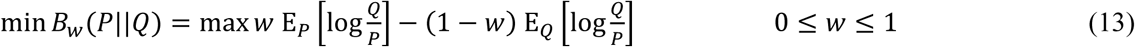

where, *w* can be tuned depending on the dropping or merging preferences. We recover *B*(*P*||*Q*), called the Jensen–Shannon divergence, when *w* is set to 0.5. Jensen–Shannon divergence is widely used in machine learning and other fields, including bioinformatics and genome analysis [28].

A closed-form expression for *B*_*w*_(*P*||*Q*) exists only when *P* and *Q* have one mixture component – e.g. the multivariate normal. In general, we should use numerical methods or an approximate solution to find the value of *B*_*w*_(*P*||*Q*), when either *P* or *Q* have more than one mixture component. Note that, the minimization problem defined in (12), which includes merging and dropping mixture components, is a combinatorial optimization problem and it becomes computationally expensive as the number of *P* mixture components grows. In the following subsections, we first derive an approximate closed-form expression for *B*(*P*||*Q*); we then propose a sub-optimal sequential merging and dropping procedure along with stopping criteria.

### 2.3. An approximate closed-form expression for *B*(*P*||*Q*)

We use a first-order Taylor expansion to approximate log *Q*(*X*)/*P*(*X*) around a point *X*_0_. We define *Q*(*X*)/*P*(*X*) as *F*(*X*), and its Taylor expansion is defined by

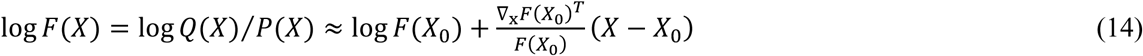

Here, we assume 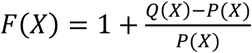 is close to 1, or 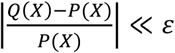. With this assumption, the first order Taylor approximation provides a good approximation of the function. We build *Q* by merging and dropping *P* components; the merging or dropping stops when the similarity between *Q* and *P* starts to significantly drop. Using the Taylor expansion defined in equation (14), *B*(*P*||*Q*) is approximated by

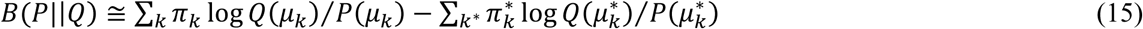

where, 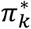 is the weight of 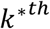 mixture component of *Q* mixture model with 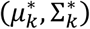 -note that, the number of mixture components in *Q* can be different from *P*. In derivation of this approximation, we linearize *F*(*X*) around the mean of each mixture component of *Q* and *P*, and then take their expectation over *P* and *Q* distributions.

Equation (15) provides a closed-form solution for *B*(*P*||*Q*). Note that, *B*(*P*||*Q*) is also a function of the mixture components’ covariances – 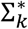 – which changes 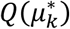 values. We can also use a second order Taylor expansion to get a better approximation of *B*(*P*||*Q*). The approximation will have a closed-form solution, which can be derived by a similar procedure described in the derivation of equation (15). Using equation (15), we can estimate the divergence measure between *P* and *Q* analytically. In both dropping and merging procedures, we examine different *Q*s built by dropping or merging *P* components and find the one with the lowest *B*(*P*||*Q*). As we discuss in the following sections, the mean and covariance of merged components are derived analytically; thus, the computational cost of the merging and dropping processes is merely the cost of computing equation (15) for different *Q*s.

### 2.4 Dropping Process

In the dropping process, we take one of the *P* components out and rescale other mixtures’ weight to keep their sum equal to one. We then check the distance between these mixture models – *Q* – and *P* to find which mixture component can be dropped. We repeat his procedure until a stopping criterion is met. The dropping process is described in Table 1:

**Table 1.**
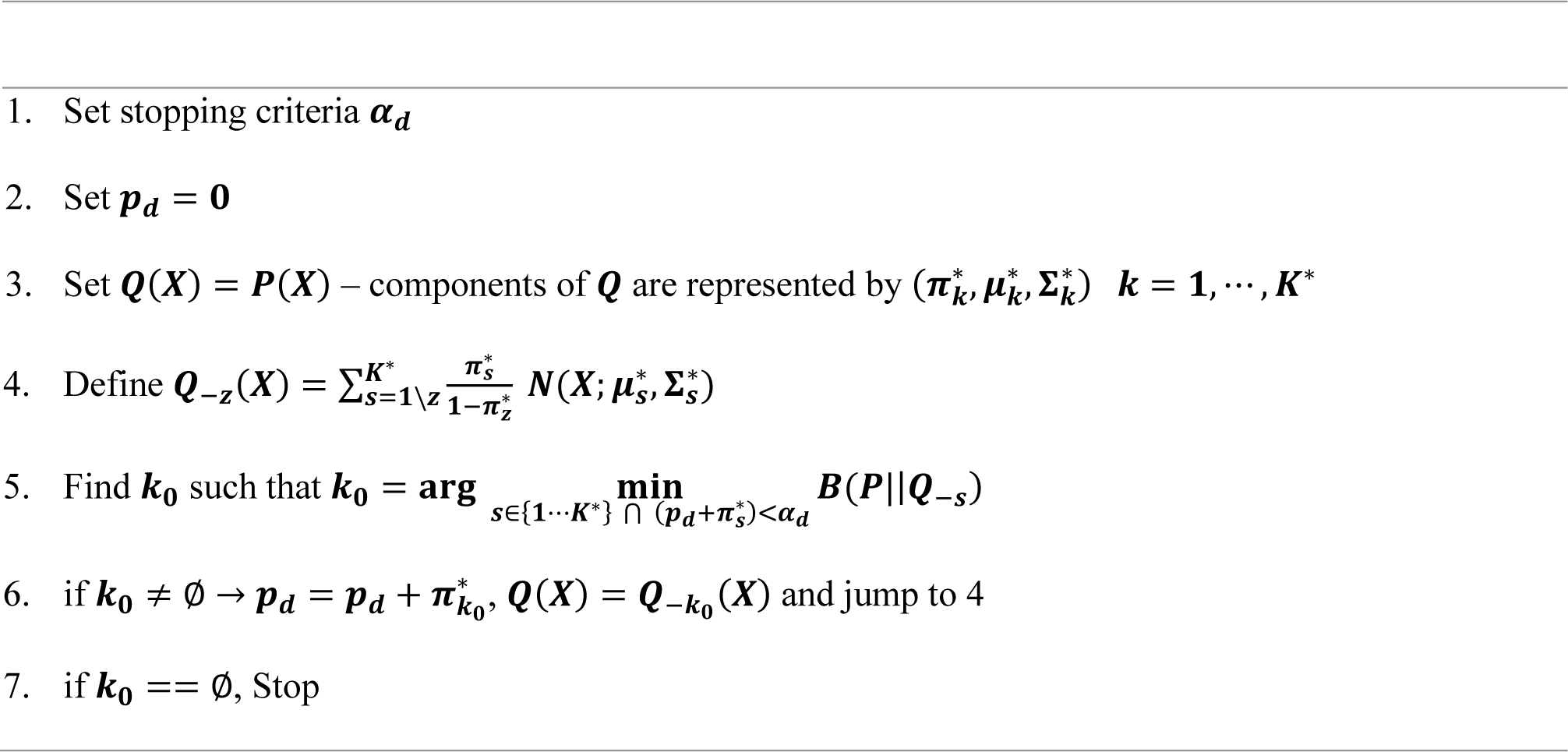
Dropping Process Algorithm

In practice, *α*_*d*_ is set to a small number – for example, 0.01. The dropping process checks which of *P* mixture components with a small mixing weight can be dropped; a component which gives the lowest divergence measure, *B*(*P*||*Q*_-*s*_). Note that the dropping process does not drop mixture components solely based on their mixing weights; this is important in optimally dropping mixture components - specifically, when there are many mixture components – or combinations of mixture components with an overall mixing weight smaller than *α*_*d*_.

We describe the analytical filter solution in Appendix B. In the filter solution, new mixture components are generated at the spike times. We first estimate parameters of these mixture components and we then call the dropping procedure to optimally drop those mixture components which contribute less in *P* distribution. The dropping process is also called on spike times; note that, the number of mixture does not grow on non-spike time steps – Appendix B.

### 2.5 Merging Process

In the merging process, we search for a pair of mixture components which can be merged while minimally increasing the divergence measure *B*(*P* || *Q*_*i*∘*j*_) – *Q*_*i*∘*j*_ represents the new mixture model with it’s *i* and *j* mixture components being merged. The merging process is run sequentially; as a result, per each iteration, number of mixture components in *Q* drops by one. The merging process is repeated until a stopping criterion is met. The merging process is defined in Table 2.

**Table 2.**
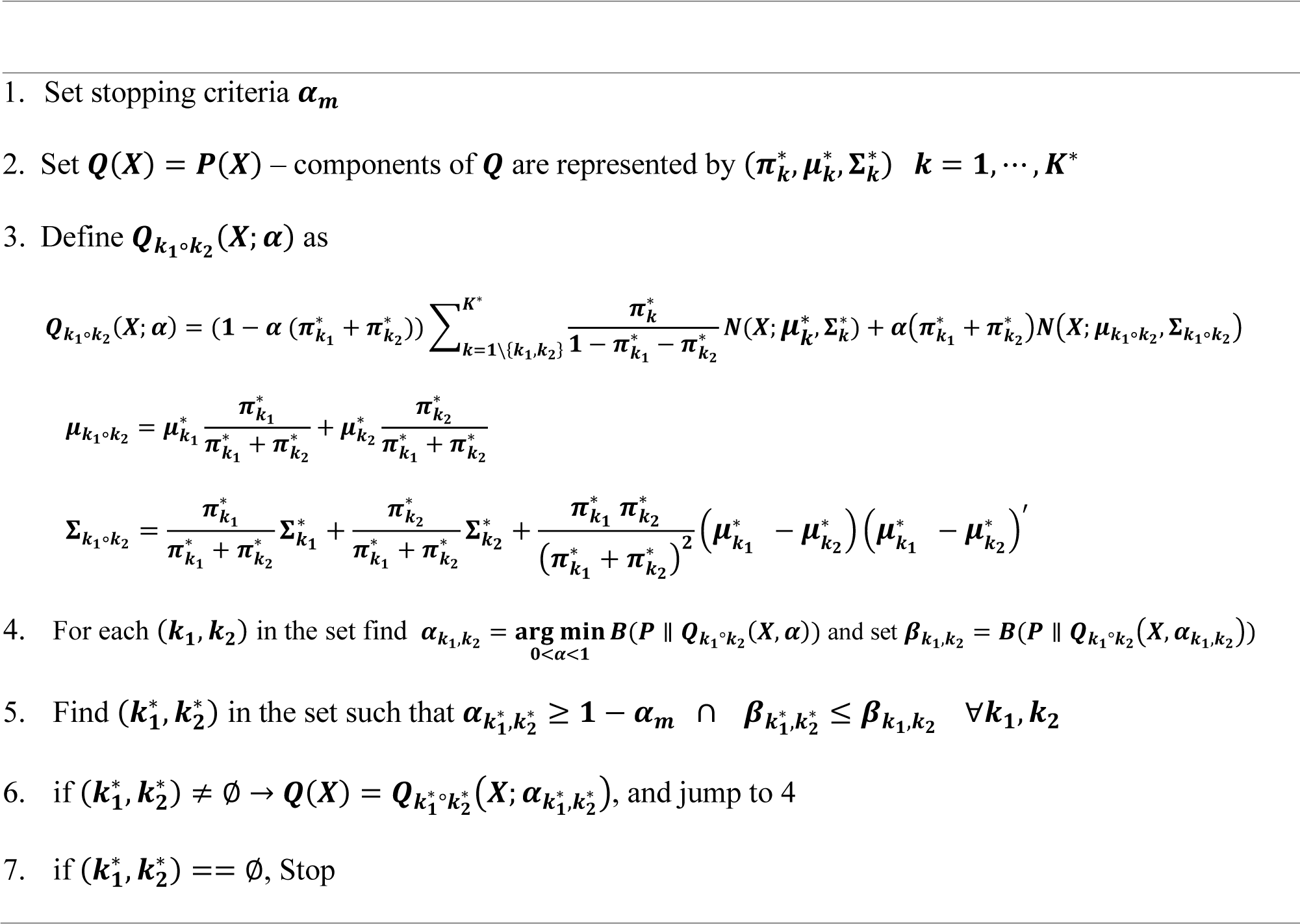
Merging Process Algorithm

In practice, *α*_*m*_ is set to a small number – for example, 0.01. For the merged components, we expect two criteria to be simultaneously satisfied. First, it must have the lowest *B*(*P* || *Q*_*i*∘*j*_); second, the merged component weight should be close to sum of two merged components weights. *α*_*m*_ checks the second criteria, and it is the largest deviation that is accepted for the discrepancy between the weight of merged component and sum of the weights of two components being merged.

The merging process is called after dropping process and it is called on each time-step. Over the non-spike periods, mixture components in the filter solution diffuse over space and this implies these components can be merged. On the spike times, new mixture components are generated by multiplying one-step prediction mixture components and the mixture components from the likelihood function. A subset of these mixture components can be merged, particularly when the likelihood function and one-step prediction mixture components coincide over space, they are good candidates for a merge.

### 2.6 Logic of the Dropping Function

The main challenge in a filter problem with a multimodal likelihood function – here, defined as a mixture model – is the exponential growth of mixture components over time. To build a computationally efficient and accurate filter solution, the key is to optimally control the number of mixtures per each processing time-step. Clearly, dropping mixture components based on their mixing weights is not an optimal solution to this problem. This is because two mixture components with the same mixing weights and different covariance matrixes cannot be treated the same. Also, without a merging process, there is always a chance that the number of mixture components will explode over time. Mixture components might also be generated with similar means and covariances, and their number will grow after being generated. Our proposed methodology addresses how these mixture components can be dropped and merged while maintaining a minimum divergence between *P* and *Q*.

A key factor in our proposed method is the definition of the divergence measure, *B*(*P* || *Q*). *B*(*P* || *Q*) penalizes the merging process – and similarly dropping process – when the merged component will change the domain of space covered by both *Q* and *P* mixture components. This is in contrast with commonly used divergence measure – like standard KL divergence, which are insensitive to changes beyond the domain of space supported by *P*. Merging only happens when a reciprocal similarity is maintained between *P* and *Q*. Thus, *Q* cannot significantly change the domain of space covered by *P* and this will lead to a more accurate approximation of *P*.

*B*(*P* || *Q*) – in its exact expression – is always positive with a minimum of zero. This implies that a part of information will be lost in the merging or dropping process, and the extent of the information loss leads to either a variance or bias error. The algorithm has multiple control mechanisms to maintain an accurate estimation of *P* over different filtering time-steps. We study different properties of drop-merge method in detail in the application section.

### 2.7 Failure Modes

The main challenge in the proposed methodology is the computational cost of the two optimization processes – and particularly the merging process – when the number of mixture components is relatively high. The number of searches over mixture pairs – plus *α* – becomes of order *o*(*K*^3^*S*) – *K* is the number of mixture components in *P* and *S* is the number of samples over *α*. In practice, we use about 10 samples over *α*; thus, a relatively large number of mixtures – for instance, larger than 100 – can lead to an expensive computation. In practice, the number of mixtures is reasonably low given that many of the mixture components are withdrawn in the dropping step.

The approximate cost function defined in equation (15) provides an analytical solution for *B*(*P* || *Q*). The accuracy of *B*(*P* || *Q*) calculation can be improved by using a second order Taylor expansion or increasing number of samples to measure *B*(*P* || *Q*) using a Monte Carlo simulation [37]. The other possible solution is to find an upper bound for *B*(*P* || *Q*) and minimize that [26]. Though, there are multiple upper bound derivation for KL distance, finding similar bounds for *B*(*P* || *Q*) is not easy. In practice, the approximation defined in equation (15) provide a reasonable approximation for *B*(*P* || *Q*) and adding more computation load for a better approximation of *B*(*P* || *Q*) might not be needed.

## 3. Application

In this section, we applied the proposed methodology to experimental data recorded by a multi-electrode array in the hippocampus of a rat. The data used in this analysis were recorded from 9 tetrodes in the CA1 and CA2 regions of the hippocampus. Spikes were detected offline by choosing events whose peak-to-peak amplitudes were above a 100uV threshold in at least one of the channels. We demonstrate the computational time and accuracy of the exact solution and the drop-merge method. For the drop-merge method, we show the result for different ranges of *α*_*m*_ and *α*_*d*_. Besides the performance and computational time, we study how the number of mixture components evolve on each processing time-step. We decode the movement trajectory in the W-maze using 2 different approaches. First, we represent the position of the maze in 1-D by only considering the linear distance of the rat from the home well, using a MoGs conditional intensity derived using the algorithm described in Ken et al. [38]. We then decode directly the rat position in 2-D, once again using a MoGs conditional intensity derived using the algorithm described in [38].

### 3.1. Decoding maze trajectory in 1-D representation

In this problem, the rat moves from the home well – **figure 1(a)** – to either the left or right arms, and it gets back to its starting point. We use a linearization scheme to express the position in 1-D by mapping the constrained linear distance from the home well to the interval [-6, 6]. In this representation, there is no distinction between left and right arms. For this problem, we used neural activity recorded from one tetrode out of 9 – implanted in the rat hippocampal area [11]; the conditional intensity mixture model built using this data consists of 35 mixture components. The update time resolution is 1 millisecond, and the state transition process variance is set at 0.01 – the variance of the state transition model is numerically estimated using the rat position data. We assume the rat movement trajectory follows a random walk model and thus we set *A*_*k*_ to 1 and *B*_*k*_ to 0. We run the model over 26 seconds of the data – about 26000 time points. **Figure 1(b)** shows the time and position of occurrence of unsorted spikes whilst the rat traversed the maze. For the exact solution, we use Riemann Sum integral method [39] with 961 samples in the range of −12 to 12 – or a 0.025 spatial resolution – to calculate the likelihood function and rat position posterior estimation. We calculate the root-mean-squared error (RMSE) between actual rat position and the mean of posterior distribution plus 95% highest probability coverage area (HPD) [40] to assess both the exact and proposed model performance. **Figure 2** shows the decoding result of the exact solution and the proposed methodology with *α*_*d*_ = 0.15 and *α*_*m*_ =0.12.

**Figure 1.**
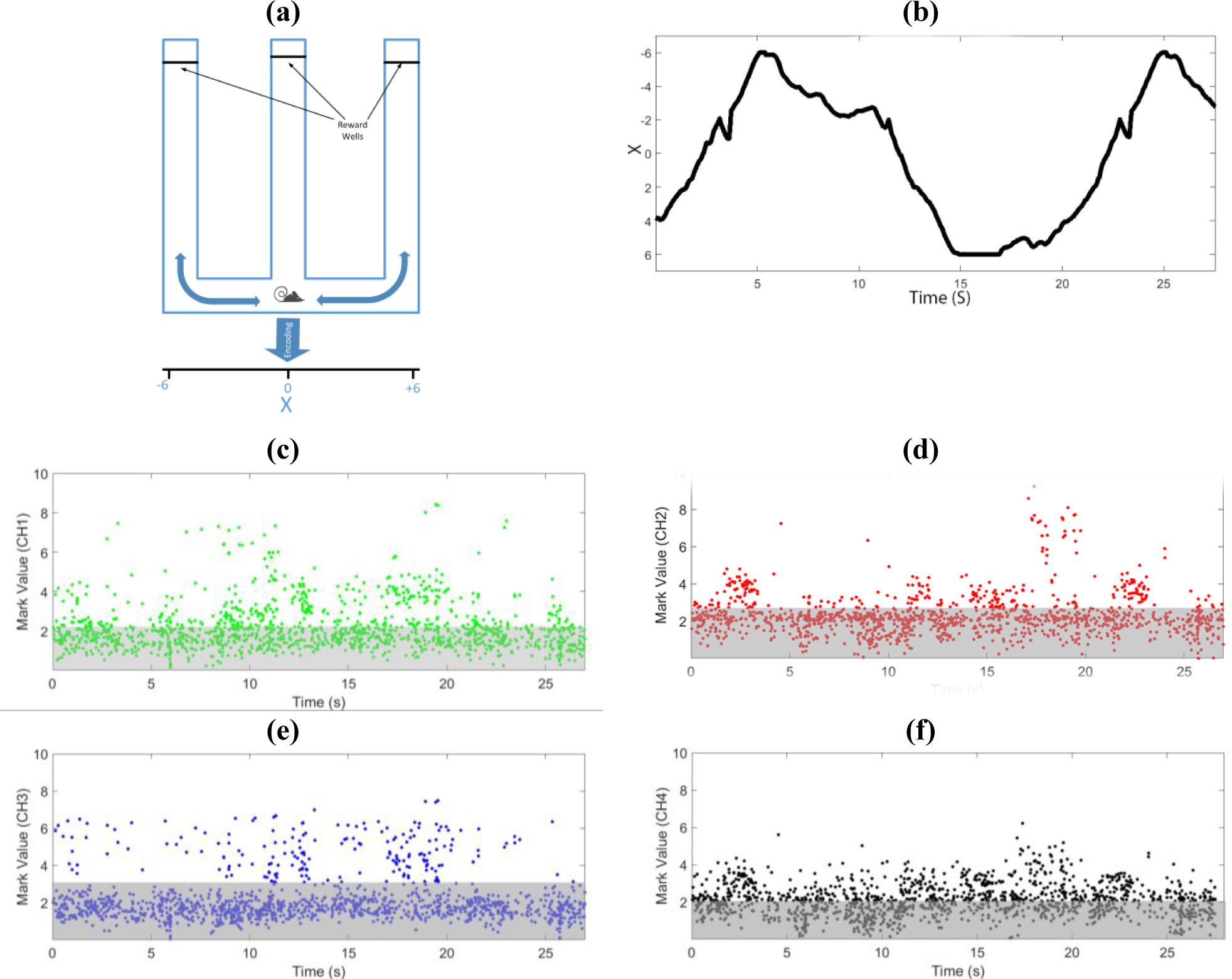
The maze structure, the rat movement trajectory, and observed neural data. **a.** We use a linearization scheme to map the 2-D position in the maze to 1-D, by mapping the constrained linear distance from the home well, to the interval [-6, 6]. In this representation, there is no distinction between left and right arms. b. Recorded Movement trajectory during 1-D task. **c-f.** Timing and mark value of observed spikes from all 4 tetrode channels. The mark values above threshold area (dark area) are more informative for decoding movement trajectory than others inside this area. Each data point present spike event. The decoding result using the drop-merge method shows a similar decoding results as the exact solution (**figure 2(a) and 2(b)**). To better assess the decoding result and computational efficiency using the drop-merge algorithm, we ran the algorithm for a range of *α*_*d*_ and *α*_*m*_ values. **Figure 3** shows the performance result and different statistics of computational time efficiency of the drop-merge method for a range of *α*_*d*_ and *α*_*m*_.

**Figure 2.**
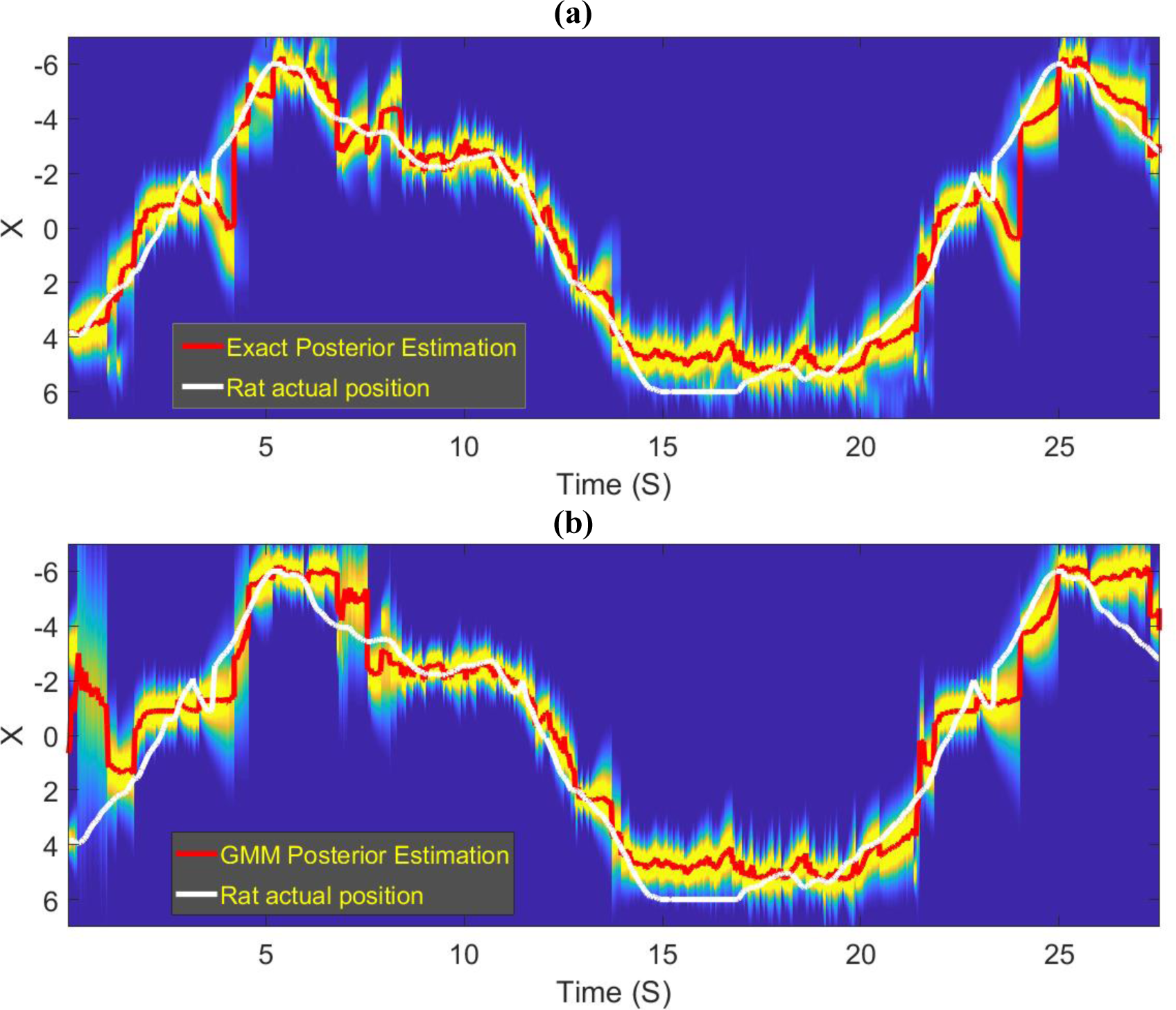
Decoding result using **a.** The exact solution and **b.** The drop-merge method with *α*_*d*_ = 0.15 and *α*_*m*_ = 0.12. Decoding result using the proposed methodology is similar to the exact solution for the most of time steps.

The processing time in the drop-merge method is variable, and it increases when a larger number of mixture components is needed to approximate the posterior distribution of the rat position. For the time steps when the number of mixture components on the previous time step is low – generally, 1, the drop-merge method runs about 200 times faster than the exact solution (**figure 3(d)**). However, when the number of mixture components on the previous time step is large, processing time of the drop-merge method becomes longer than the exact solution. For example, for *α*_*m*_ and *α*_*d*_ equal to 0.1, there are time steps with a processing time 2.6 times larger the average processing time in the exact solution (**figure 3(e)**). However, this situation is the worst scenario and it only happened in less than 0.17% of time steps in our example.

**Figure 3.**
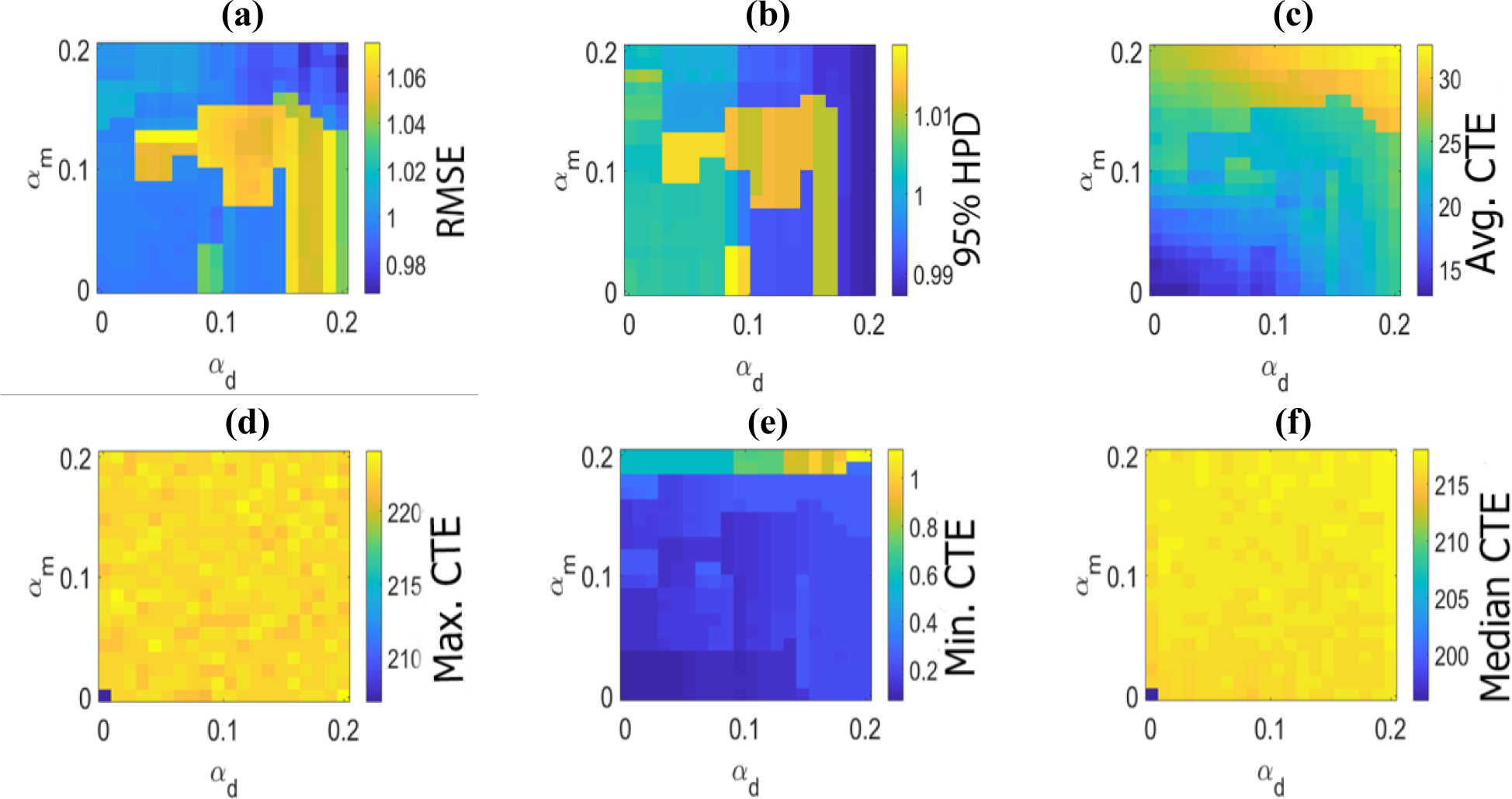
Performance and computational saving of drop-merge algorithm for 1-D decoding task using different *α*_*d*_ and *α*_*m*_parameters - *α*_*d*_ and *α*_*m*_ are the drop and merge stopping criteria defined in Table 1 and 2. Each performance map in **a**, **b** is normalized to the corresponding result derived from the exact filter solution. **a.** RMSE performance map. Value of 1 corresponds to a similar RMSE measure for the exact and proposed method. A lower value reflects more accurate decoding. **b.** 95% HPD coverage performance map. A value close to 1 corresponds to a similar coverage area for both the exact and proposed method. A larger value is more desired. **c**. Average computational time efficiency (CTE) using the drop-merge method. In **c** to **f** figures, the average processing time in the exact method is divided by the average processing time per time step using drop-merge method. For the exact solution, we use a Riemann Sum integral with a 0.025 step. The 0.025 is the coarsest resolution which maintains the exact solution’s performance, when it is run with much finer resolutions. A larger value is more desired, for instance, a value of 20 implies that the drop-merge method run 20 times faster than the exact solution **d.** Maximum computational time efficiency using the drop-merge method. For instance, a value of 215 implies that for the corresponding parameter setting there is at least one time step where the computation saving is 215 times faster than average processing time of the exact solution. **e.** Minimum computational time efficiency using the drop-merge method. For instance, a value of 0.5 implies that for the corresponding parameter setting, the longest processing time of the drop-merge method is twice the average processing time in the exact solution. **f.** Median of computational time efficiency using the drop-merge method. For instance, a value of 210 implies that for the corresponding parameters setting, half of time steps run at least 210 faster than exact solution.

Note that per each time-step, we can change *α*_*d*_ and *α*_*m*_ depending on the number of mixture components needed to be processed avoiding the instances with a long processing time. The average processing time in the drop-merge method is 15 times faster than the exact solution independent of choice of *α*_*m*_ and *α*_*d*_values. By defining an optimal choice for *α*_*d*_ and *α*_*m*_, we can even gain a higher computational time efficiency. In Appendix D, we show the histogram of processing time for the exact solution and drop-merge method with *α*_*d*_ = 0.15 and *α*_*m*_ = 0.12 to provide a better picture about different methods’ the processing time statistics.

Using the performance result presented in **figure 3**, we can choose specific values for *α*_*d*_ and *α*_*m*_ based on our goals for speed and accuracy. Here, we picked *α*_*d*_ =0.15 and *α*_*m*_ = 0.12 for the drop-merge method, which gives about 22 times faster processing time than the exact solution. The RMSE of the drop-merge method is only 6% higher than the exact solution, and the 95% HPD is almost the same as the exact solution. Table 1 provides further information on the drop-merge method for this choice of *α*_*d*_ and *α*_*m*_.

**Table 1.** Performance result using the exact and proposed solution in 2D decoding.

| Method | Setting | RMSE (cm) | 95% HPD | Processing Time (msec) | Avg. Number of Mixtures | Average Number of Mixtures on Spike Time | Maximum Number of Mixtures |
| --- | --- | --- | --- | --- | --- | --- | --- |
| Exact Solution | $dx = 0.025$ | 0.80 | 80.5 | 24.4 | NA | NA | NA |
| GMM-based method | $\alpha_d = 0.15$<br>$\alpha_m = 0.12$ | 0.85 | 81.0 | 1.1 | 1.21 | 2.8 | 11 |

The comparison results show that the drop-merge algorithm is capable of reducing the decoding computational time, which is the main goal of this algorithm. Note that the computational time efficiency is attained without substantial decreases in accuracy. The result reported here demonstrates the algorithm potential in tracing the exact solution, whilst saving the computational cost; the computational efficiency becomes a more critical factor in high-dimensional decoding problems. Thus, now we study properties of the drop-merge algorithm in a 2-D decoding problem.

### 3.2 Decoding maze trajectory in 2-D

The previous section projected a 2-D position onto a 1-D representation and ignored the distinction between left and right. In reality, the animals horizontal position is a 2-D variable, and here we develop a solution for this more realistic case. While in the 1-D decoding problem, we used neural recording from one tetrode or, just one cells ensemble – in building the conditional intensity and decoding step; here, we use the recordings from nine different tetrodes in both the encoding and decoding step. We find that with six or more tetrodes, we observe a satisfactory decoding result; however, we use a larger number of tetrodes to better assess the performance and computational cost of the exact and drop-merge method. We build the conditional intensity for each tetrode individually and then use them together in the decoding step – see Appendix A for a description of extending the encoder and decoder model to the case where there are conditional intensities from multiple tetrodes. **Figure 4(a)** shows the movement trajectory in 2-D space; here, each point of the maze - or, the rat position - has a distinct coordinate.

**Figure 4.**
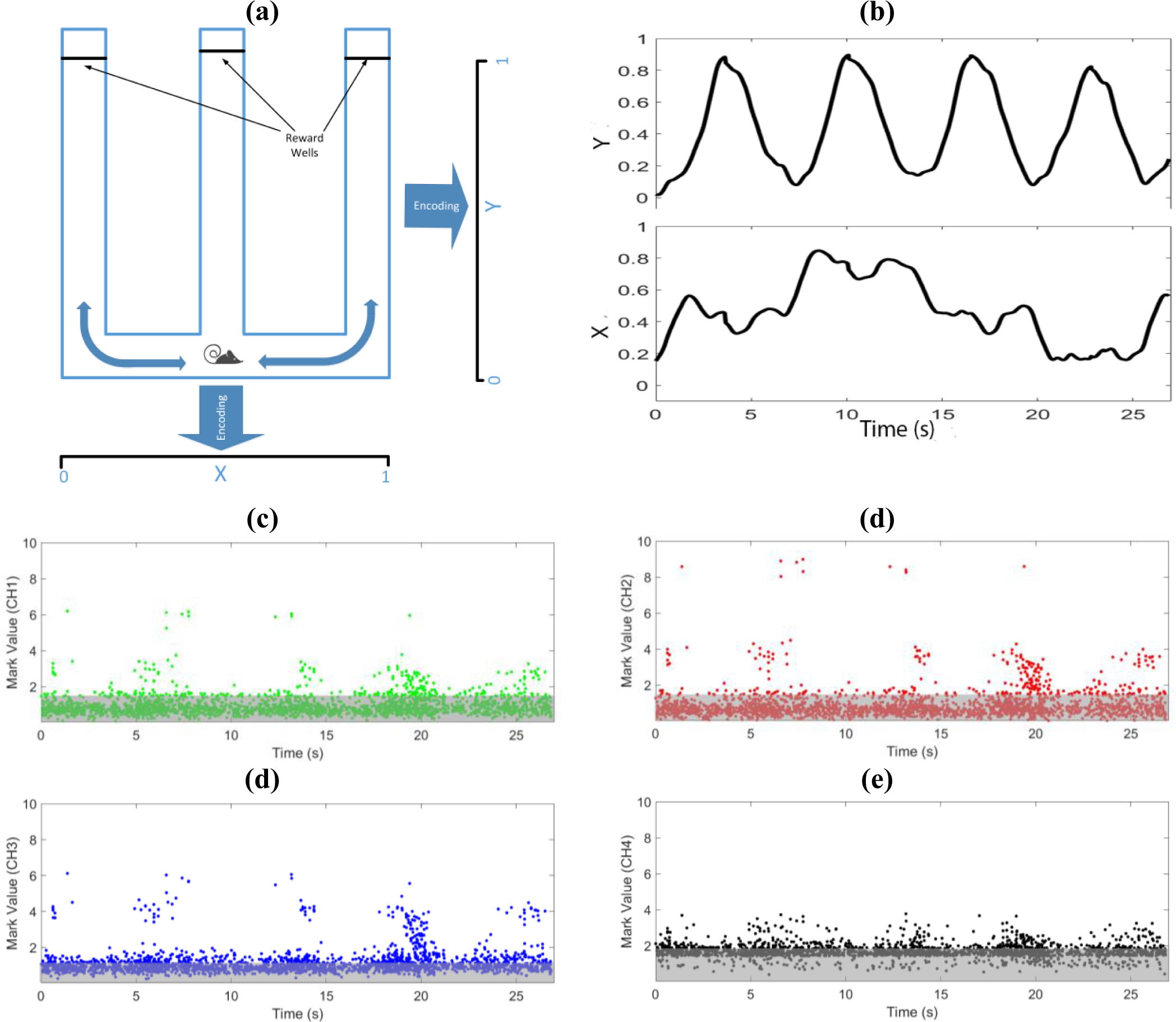
The maze structure, the rat movement trajectory, and sample neural data. **a.** W-maze, the rat moves from the center arm to the left and right arms to get food reward. The rat coordinates are scaled from 0 and 1 – in the figure, (0.5, 0) is the coordinate of the rat position shown in the figure. **b.** Both movement trajectories along X and Y directions. **c-f** Timing and mark value of observed spikes from all channels for one of tetrodes. The mark values above threshold area (dark area) are more informative for decoding movement trajectory than others inside this area. For the exact solution, we use Riemann Sum integral method [39] with 300 samples in the range of −1 to 2 – corresponding to 0.01 spatial resolution – to calculate the likelihood function and rat position posterior estimation. The finer spatial resolution in 2-D compared to the value being used in 1-D decoding addresses the change in coordinate scale used to represent the rat position in the maze. We calculate the same performance measures as we used in the previous section to assess both the exact and drop-merge model performance.

**Figure 5** shows the decoding result of the exact solution and drop-merge methodology at two different points with two different sets of parameters. For *α*_*m*_ = 0.15 and *α*_*d*_ = 0.15, the number of mixture components for the first and second time steps are 3 and 5, while for *α*_*m*_ = 0.1 *and α*_*d*_ = 0.05, the number of mixture components for the first and second time steps are 5 and 5. The decoding result using the proposed methodology shows a similar posterior estimate of the exact solution in these sample time points.

**Figure 5.**
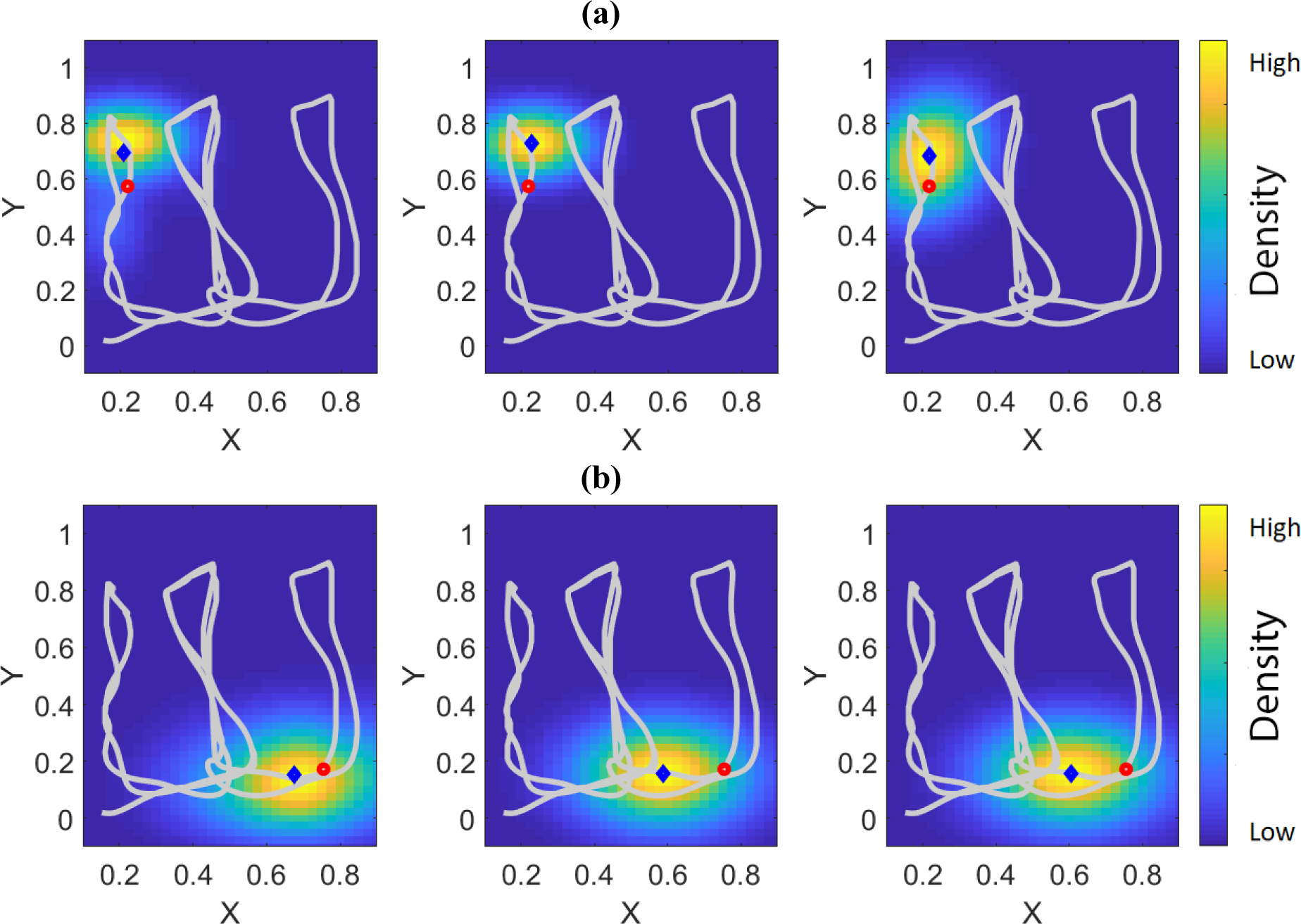
Decoding result for the exact solution and drop-merge method at two different time steps using two different merge and drop parameter settings. The left panel shows the decoding result using the exact method. The middle panel is the merge-drop method decoding result with α_d_ = 0.15 and α_m_ = 0.15 and the right panel shows the decoding result with α_d_ = 0.1 and α_m_ = 0.05. **a**. Decoding result on the first sample point. **b**. Decoding result for the second sample point. In these figures, Red dot shows the rat actual position and blue dots shows the mean of decoder posterior estimation.

To better assess the decoding result and computational time efficiency using the drop-merge method, we ran the drop-merge method for a range of *α*_*d*_ and *α*_*m*_ values. **Figure 6** shows the performance result and different statistics of computational time efficiency of drop-merge method for a range of *α*_*d*_ and *α*_*m*_.

**Figure 6.**
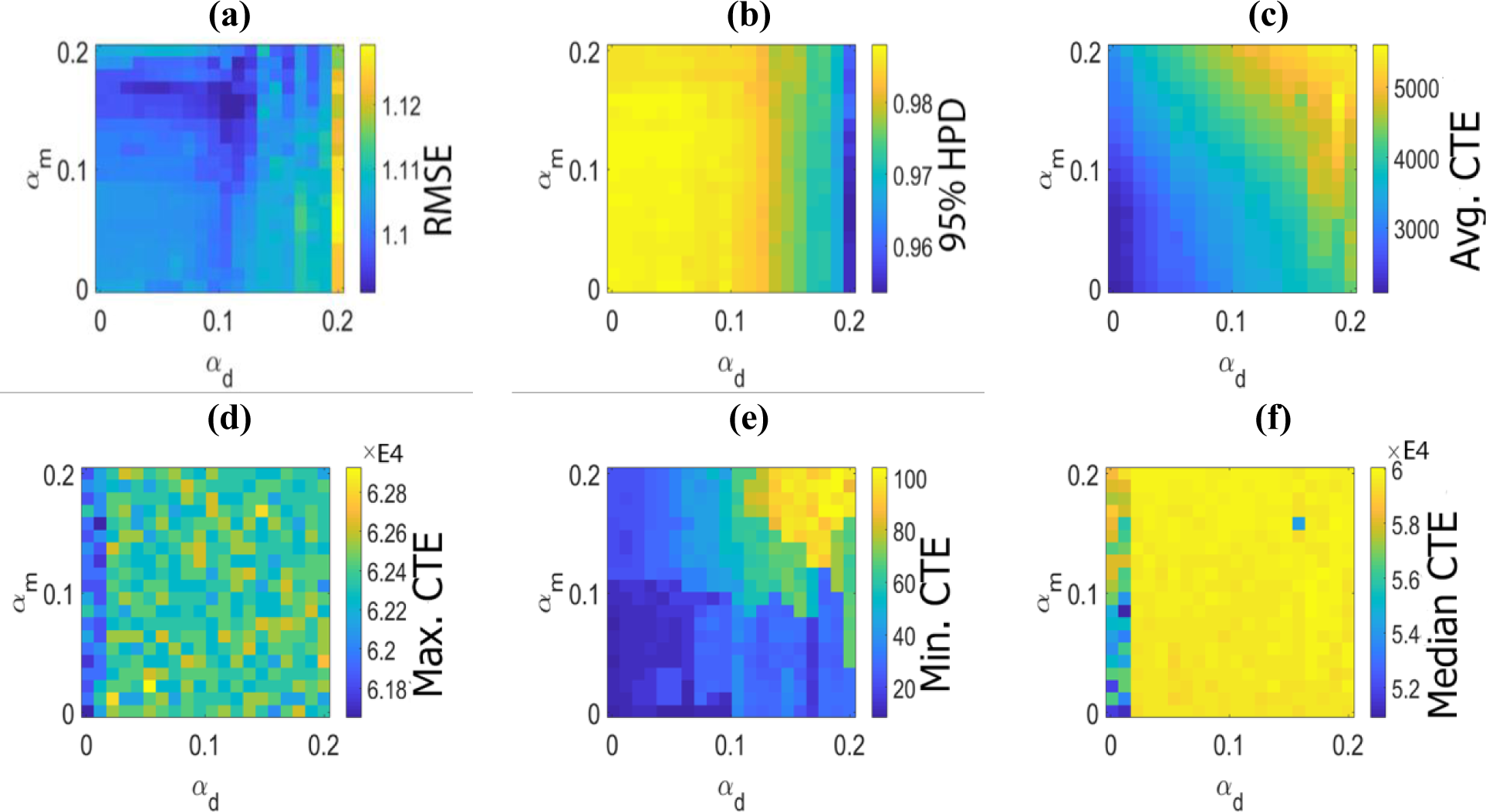
Performance and computational time efficiency of the drop-merge method for 2-D decoding task using different *α*_*d*_ *α*_*m*_ and *α*_*m*_ parameters. Each performance map is normalized to the corresponding result derived from the exact filter solution. **a.** RMSE performance map. Value of 1 corresponds to a similar RMSE measure for the exact and proposed method. A lower value is more desired **b.** 95% HPD coverage performance map. A value close to 1 corresponds to similar coverage area for both the exact and proposed method. A larger value is more desired. **c**. Average computational time efficiency (CTE) using the drop-merge method. Here, the average processing time in the exact method is divided by the average processing time per time step using the drop-merge algorithm. For the exact solution, we use a Reimann Sum integral with a 0.01 spatial resolution. A larger value is more desired; for instance, a value of 4000 implies that the drop-merge method run 4000 times faster than the exact solution **d.** Maximum computational time efficiency using the drop-merge method. A value of 62000 implies that the for corresponding sets of parameters, there are time steps that runs about 62000 times faster than the exact solution. **e.** Minimum computational time efficiency using the drop-merge method. **f.** Median computational time efficiency using the drop-merge method.

**Table 2** shows the performance of the exact and proposed methodology in the 2-D decoding for *α*_*d*_ = 0.1 and *α*_*m*_ = 0.05. The computation time of both methodologies are reported as well.

**Table 2.**
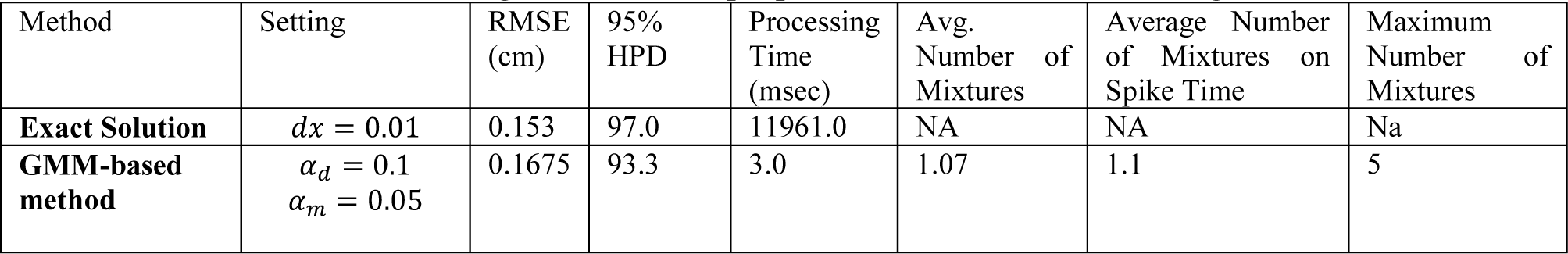
Performance result using the exact and proposed solution in 1-D decoding problem

| Method | Setting | RMSE (cm) | 95% HPD | Processing Time (msec) | Avg. Number of Mixtures | Average Number of Mixtures on Spike Time | Maximum Number of Mixtures |
| --- | --- | --- | --- | --- | --- | --- | --- |
| **Exact Solution** | $dx = 0.01$ | 0.153 | 97.0 | 11961.0 | NA | NA | Na |
| **GMM-based method** | $\alpha_d = 0.1$<br>$\alpha_m = 0.05$ | 0.1675 | 93.3 | 3.0 | 1.07 | 1.1 | 5 |

The performance result and computational time efficiency in the 2-D decoding problem are aligned with the result observed in 1-D decoding problem. We get even more computational time efficiency with our proposed method, as desired. The performance result of the drop-merge method is similar to the exact method, while its computation time is at least 2500 times faster than the exact method. The average processing time is about 3.0 milliseconds for each time step – 1 millisecond time interval. Here, we implemented these algorithms in MATLAB (MATLAB Release 2017a The MathWorks, Inc., Natick, Massachusetts, United States) platform and scaling their computational time by 3 or 4 times to get below 1 millisecond should be achievable by optimizing the code or implementing it using Matlab Mex Compiler, Python, or C++.

## 4. Discussion

We developed an approximate filter solution for a class of marked point-process filter problems, in which the conditional intensity of the neural activity of an ensemble of cells, is defined by a MoGs. In developing the solution, we approximated the posterior distribution using a GMM. We then proposed a drop-merge method, which collectively estimates the optimal number of mixture components plus their corresponding parameters - weight, mean, and covariance matrix - per each time-step. We further examined drop-merge algorithm in both 1-D and 2-D decoding problems using neural data recoded from rat hippocampus place cells. For 1-D problem, we use neural recording of one tetrode to estimate the rat position. In 2-D decoding problem, we applied the drop-merge method for multiple tetrodes recording which is a more realistic scenario for a decoder problem. The result in both problems show the capability of the proposed method in conjunction with GMM encoder model for a real-time neural decoding in 2-D and even higher dimensional decoding problems.

The specific aim of this research was to develop a computationally efficient decoder model. This requires building a concise picture of which factors will affect the merge and drop processing time. The drop and merge algorithms are combinatorial optimization problems; thus, their associated computational time is determined by the initial number of mixture components being processed. The dropping and merging algorithms iteratively drop the mixture components until the corresponding stopping criterion is met. For the initial mixture model *P* with *K* mixture components, the maximum number of iterations for dropping or merging algorithms is *K*. In the *m*^*th*^-iteration of the dropping or merging algorithm, we compare *K* - *m* - 1 different mixture models in the dropping algorithm, or (*K* - *m*) × (*K* - *m* - 1)/2 different mixture models in the merging algorithm to find which of these new mixture models with either dropped or merged components give the lowest divergence measure. The dropping algorithm is computationally less expensive than the merging algorithm; thus, it is called first to reduce the number of mixture components being passed to the merging algorithm. When the expected number of mixture components are limited, the merging algorithm can be called without the dropping algorithm. Each internal iteration of the drop or merge itself becomes more expensive with an increase in *K*, as the cost of calculation of *P* and *Q* in the divergence measure increases with *K*. *P*(*X*) and *Q*(*X*) must be evaluated at *K* different values of *X*, and for *KL*(*Q*||*P*), we calculate these values for *K* - *m* different points of *X*, because *K* - *m* is the number of mixtures in *Q* after the *m*^*th*^ call of either merging or dropping algorithms. However, we can reduce this computational load using values of *P*(*X*) and *Q*(*X*) calculated in the previous iteration of the merging and dropping algorithms. Note that in the dropping algorithm, a mixture’s mean and covariance are the same in each iteration, except their mixing weights are changing on each iteration. Additionally, *KL*(*Q*||*P*) is estimated using a subset of *X* points that already calculated in the previous iteration of dropping algorithm. Thus, we can use previous values of *P*(*X*) and *Q*(*X*) – specifically, values of their mixture components – to significantly reduce the computation time of calculating *KL*(*Q*||*P*) and *KL*(*P*||*Q*), and thus the computational time of *B*(*P*||*Q*). For the merging algorithm, the mean and covariance of only one mixture component changes from one iteration to the next, while the mean and covariances of all other mixture components stay the same as previous merging iteration. Thus, we can use these values to reduce the computational time of calculating *B*(*P*||*Q*) in the current iteration of the merging algorithm. Overall, the computation time of *B*(*P*||*Q*) in the dropping algorithm is less than merging algorithm; thus, the dropping algorithm is called first as it has the lower computational cost. The computation time analysis for the more demanding 2-D decoding, shows that merging algorithm can deal with up to 10 mixture components in *P* within 1 millisecond computational time. The dropping process takes less than 1 millisecond to run on a GMM with up to 70 mixture components, and it is called ahead of merging process to bring the number of mixtures down. In general, we can run both the dropping and merging algorithm in less than 2 milliseconds per processing time-step, which is fast enough for real-time decoding application. Note that these results are based on the algorithm implementation in MATLAB – a scripting programming language, and reducing this computation time to less than 1 millisecond is feasible for structured programming platforms like C++ [41], Python (Python Software Foundation, https://www.python.org/), or MATLAB Mex Compiler (MATLAB and Build MEX Toolkit Release 2017a The MathWorks, Inc., Natick, Massachusetts, United States).

The drop and merge algorithms solely deals with the mixture components and it is independent of the processing step that generate the mixture model; thus, we can use other approximation techniques to build GMM models and utilize the merging and dropping algorithm to manage the growth in the number of mixture components over time. We use a first order Taylor expansion to find an analytical solution for *B*(*P*||*Q*) calculation. We can use other approximation techniques like the second order Taylor expansion, approximate upper bound [26], or Monte Carlo simulation [37]. Note that in both dropping and merging algorithms, we have a secondary mechanism – characterized by *α*_*d*_ and *α*_*m*_ – which limit the extent of divergence between *P*(*X*) and *Q*(*X*). Under this control mechanism, the first order Taylor expansion gives a good estimate of *B*(*P*||*Q*). However, both the merging and dropping algorithms can be run using different estimations of *B*(*P*||*Q*).

The filter solution in 1-D decoding problem is not computationally expensive; however, it becomes expensive in 2-D decoding problem and almost interactable in higher dimensions. The filter solution using GMM posterior proposed here accompanied with the merge-drop algorithm can be a proper solution for the decoding problems with higher dimensional spaces. The average computation time in drop-merge method is about 3 milliseconds for 2-D decoding, and this is about 4000 times faster than the exact solution. The computation time in 2-D decoding has only increased from 1 millisecond - in 1-D decoding - to 3 milliseconds, whilst the exact solution computation cost jumped from 24.4 milliseconds to 11909 milliseconds (about 12 seconds). For the merging and dropping algorithms, the computational cost in the algorithm arises only in calculation of *P*(*X*) and *Q*(*X*) distribution, not the algorithm itself. Thus, we expect the computation cost grow linearly by the dimension of the problem and this makes the algorithm suitable for high-dimensional decoding problem.

The specific aim of this research was to build a computationally efficient decoder model; however, proposed solutions must retain a comparable performance with regard their decoding accuracy. We studied two performance metrics – RMSE and 95% HPD – to compare the decoding result using the exact and our proposed filter solutions. For *α*_*d*_ = 0.15 and *α*_*m*_ = 0.12, RMSE is only about 6% percent above the exact solution. The 95% HPD coverage area is 81.0 percent which even 0.7% percent better that the exact solution. A similar performance trend can be seen for the 2-D decoding problem; for *α*_*d*_ = 0.1 and *α*_*m*_ = 0.05, we get about 4000 times computational efficiency compared to the exact solution. This computational saving comes with 9% increase in RMSE and with only 3.7% drop in the 95% HPD coverage area measure. The result suggests that the approximate filter solution along with the merge-drop algorithm give an accurate decoding performance whilst reaching a significant computational saving.

Though our proposed solution provides a boost in computational efficiency, a better picture of how this saving emerge from will helps us to address future improve of the framework. We examined in Appendix C, how the computational saving changes on the spike and non-spike times for merging-dropping algorithms. The computational saving in 1-D decoding mostly comes from non-spike time points, where the merging and dropping algorithms are run on GMMs with a small number of mixtures. The merging-dropping algorithm provides a parametric distribution for the rat position on each filtering time step, and thus, this makes benefiting from non-spike time in reducing computational time possible. Note that in the dataset used in this analysis – for 1-D decoding, 95% of data points are non-spike. For 2-D decoding, we observed a consistent computational saving in both spike and non-spike time points. Though the computational saving in non-spike timing is an order of magnitude larger than the spike-time, we get a significant boost in the computational time. Principally, we aim to use the proposed methodology in this research in a multi-dimensional decoding problem, and its computational benefit has been reflected in 2-D decoding studied here. Sparseness of the neural activity plays a significant rule in boosting the computational efficiency of our filter solution; in other work, we gain from our signal properties in reaching a better computational saving. This neural property is present independent of the dimension of the decoding problem, and it is retained in higher dimensional decoding problems.

A distinct component of the merge-drop algorithm proposed here is its divergence or distance metrics. We used a symmetric divergence measure in our merging and dropping algorithms, and we argued that the choice of divergence measure is an important factor in reaching a comparable performance to the exact method. In Appendix C, we run the same 1-D decoding problem described in the application section using our proposed algorithms with a KL divergence measure. Using KL divergence, the performance metrics degrade significantly. Also, the average number of mixtures per processing time-step is lower than the symmetric measure. The KL divergence measure tends to merge mixture components, and this leads to a lower number of mixtures in average. Though computation wise this is a desired phenomenon, this leads to a biased posterior distribution of the rat position, degrading the overall performance in drop-merge method using a KL distance measure.

The merge-drop algorithms proposed here are not parameter-free models; however, we only need to pick two parameters, one for the merge and one for the drop algorithm. For the examples demonstrated here, we picked optimal *α*_*d*_ and *α*_*m*_ by considering a balanced performance and computational time saving. Depending the decoding problem objective, we can pick different values of *α*_*d*_ and *α*_*m*_. By selecting smaller *α*_*d*_ and *α*_*m*_ parameters, we get almost the same performance of the exact solution – **figure 3(a-b)** and **figure 6(a-b)**, while the algorithm is an order of magnitude faster than the exact solution. To get even higher computational saving, we can change *α*_*d*_ and *α*_*m*_ on every processing time step given the number of mixture needs to be processed. Thus, the drop-merge method is suitable for almost all decoding problems in hand by changing its *α*_*d*_ and *α*_*m*_ parameters properly.

The approximation filter solution using GMM and a MoGs conditional intensity can be applied for other filter problems, where the posterior has a complex and multi-modal distribution. Mixture models are powerful and flexible tools to approximate complex and multi-modal functions and distributions like modeling neural activity in response to external stimuli. The idea of using mixture models to characterize neural activity and utilize them in developing computationally efficient inference steps like filter has a great promise in the neural decoding problems. The importance of mixture models becomes even more significant when a filter solution needs to be developed for multi-dimensional decoding problems.

So far, we discussed computational saving and performance of the approximate filter solution and the merge-drop algorithms. However, there are other steps need to be taken or addressed in future research to further enhance accuracy and particularly computational efficiency of the proposed solution. Though the exact solution becomes computationally expensive, its processing time per each processing step is predictable. In contrast, the processing time in the merge-drop solution is a function of the number of mixture components being passed to these algorithms. When the number of mixture components become large, these algorithms require a large time to compare different pairs and this causes processing time to be long. We need to find the solution like optimal choices for *α*_*d*_ and *α*_*m*_ to control the overall computational time per processing time or identifying the candidate mixture components which will be merged or dropped in place of checking all possible pairs. The proposed merge-drop solution is being built upon one-step optimality; in other words, we assume the approximate solution on each point is an accurate representation of the filter solution, and we build the next time step filter solution based on this assumption. However, a more accurate solution requires accounting the performance accuracy on the next processing steps as well. The idea is how we can use the sparseness property of neural activity to build a more accurate filter solution. For example, we should consider that each spiking activity follows by a non-spiking period and build the approximate filter solution which optimizes the filter solution on the current and future processing time. Here, we build a partial two-steps optimality and how we can extend this optimality to longer period is of a great interest.

## 5. Conclusion

In this article, we proposed a computationally efficient filter solution for the marked point-process filter, when the observation conditional intensities (CIFs) are defined by a MoGs. In developing this solution, we assumed the posterior distribution at each time step can be approximated by a GMM, and we derived the solution to optimally define the number of mixture components and their corresponding parameters (mixing weights, mean, and covariance) for each filtering time step. For a conditional intensity function defined by a MoGs and random-walk state transition process, we show the posterior estimation can be reasonably approximated using a new GMM. Under this assumption of posterior and conditional intensity, the number of posterior mixture components grows each time step, and the merging and dropping procedures proposed here provides a systematic procedure to optimally control the number of mixtures, while it maintains a pre-defined level of similarity to the exact solution. We demonstrated the solution in both 1-D and 2-D decoding probelms, and we showed that our proposed solution maintains a similar performance to the exact solution, while its computational time is significantly lower. The computational cost drops below the data update time, which makes the solution suitable for real-time applications. The proposed methodology can be applied to other non-linear or higher-dimensional filter problem, where an accurate solution with a minimal computational time is needed.

## Acknowledgments

This research was partially funded by R01 MH105174 and SCGB grant #320135. A.Y. and M.R.R. wrote the manuscript with support from K.A. and U.T.E. A.Y. developed the theory and algorithm implementation. A.Y. and M.R.R. performed data analysis. We would like to thank Dr. Eric Denovellis for implementing the algorithm in Python and debugging the source code.

A copy of the source code utilized in this research along with sample data can be found in the following GitHub link: https://github.com/Eden-Kramer-Lab/GMM_PointProcess

## Appendix A Likelihood Function for Activity of Multiple Cell Ensembles

Let’s assume we have C independent extracellular electrodes, each one is recording the spiking signals from an ensemble of neurons. In the interval Δ_*k*_, we observe 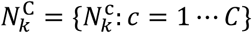 events, where 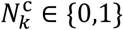 and it defines spiking event of *c*^*th*^ ensemble of cells’ neural activity with 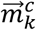 mark - 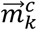 is observed when 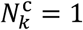. We can assume mark spike events of each of these C cell ensembles are independent of each other given the history term and external covariate - *X*_*k*_; thus, the likelihood of *X*_*k*_ given the spike mark events is defined by

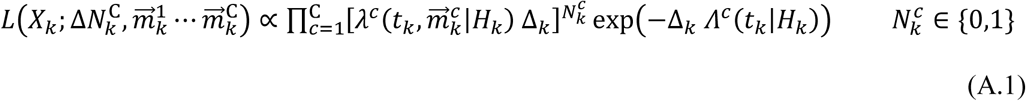

Note that the exponential term - exp(−Δ_*k*_ *Λ*^*c*^(*t*_*k*_|*H*_*k*_)) – is present in the likelihood function independent of observing a spike or not. Let’s define

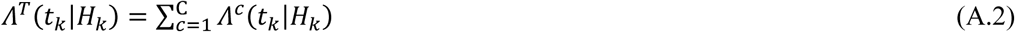

We can also define sum of observed event in Δ_*k*_ by

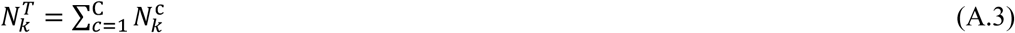

When 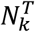 is zero, the likelihood function is only defined by *Λ*^*T*^(*t*_*k*_|*H*_*k*_). However, when 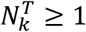, we can partition Δ_k_ to smaller time intervals defined by 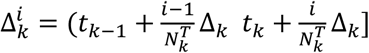 and assign one of mark spike events to each of 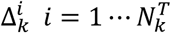. Let’s assume c_i_ is one of those ensembles with a spike event, then the likelihood function for 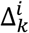 is defined by

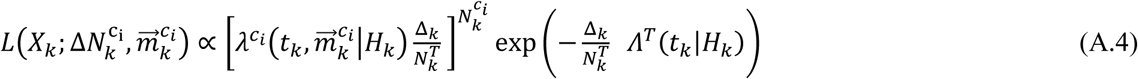

Here, we factorize the likelihood function to multiple likelihood function calculated in shorter time intervals. We can also properly scale ∑_*Q*_ – random walk covariance matrix – 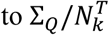 and run one-step prediction and update rule in this partitioned time intervals. Under this modeling assumption, the likelihood function in equation (A.4) is the same as equation (5) defined when we only have the joint mark conditional intensity and ground conditional intensity for the activity of one cell ensemble. In (A.4) definition, we ignored different time sequence of those 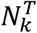 events in Δ_*k*_ interval; thus, we can have 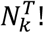 different combinations in building the filter solution when there is more than one spike mark event in an interval. Under this modeling assumption, we change the interval of filter update given the number of events being observed in a time interval. We could also start with shorter time interval for each time step, where the probability of observing more than one event in the interval is infinitesimal.

## Appendix B Closed Form Solution for Posterior Distribution of State – *X*_*k*|*k*_ – Under Assumption of a Mixture of Gaussians (MoG) for the Join Mark Intensity Function

Let’s assume the posterior distribution of state at time *k* - 1 (*X*_*k*-1|*k*-1_) is defined by equation (8.a).

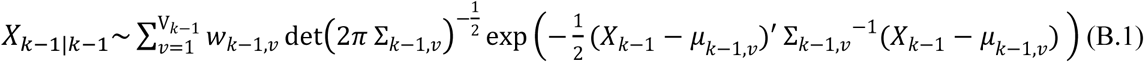

where, V_*k*-1_ defines number of mixtures and (*μ*_*k*-1,*v*_, ∑_*k*-1,*v*_) are the mixtures’ mean and covariance estimates. The one-step prediction – defined in equation (8.a) – for the state transition process defined in equation (7) can be described by

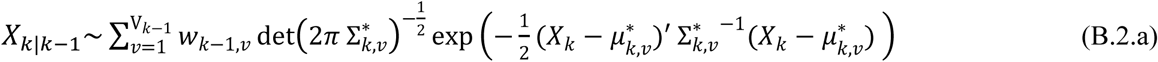

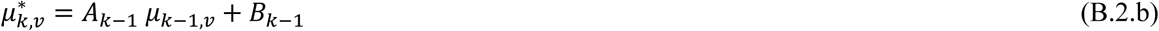

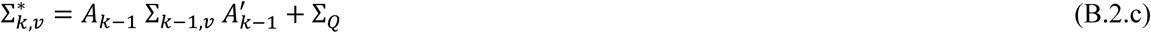

Let’s assume there is a spike with mark 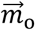 at time *k*. By replacing 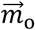 in equation (2), we have

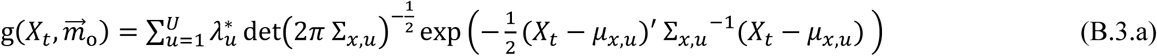

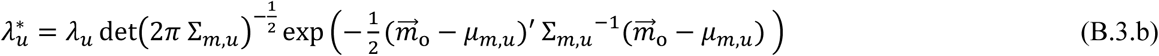

where, 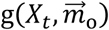 becomes a MoG with known – unnormalized - weights.

For the filter update on a spike time, 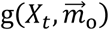 is multiplied by *X*_*k*|*k*-1_ – we call this new term 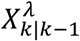. There are *U* mixture components in 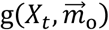 and V_*k*-1_ mixtures in *X*_*k*|*k*-1_; their multiplication will generate a new MoGs with V_*k*-1_ U mixture components. This is because multiplication of two multivariate normal distributions with (*m*_1_, ∑_1_) and (*m*_2_, ∑_2_) is a new multivariate normal [42] defined by

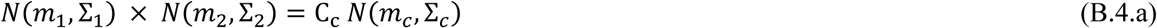

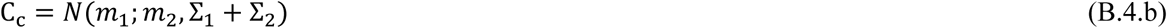

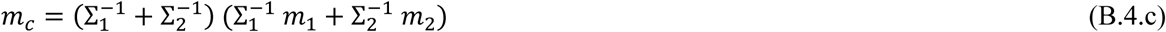

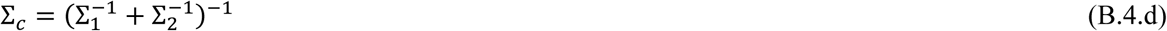

where, (*m*_*c*_, ∑_*c*_) define the new components’ mean and covariance and C_c_ is a scaling term – if the *w*_1_ and *w*_2_ are the weight of those two components, the new component weight becomes *w*_1_ *w*_2_ C_c_.

The g(·,·) term only appears on spike times; thus, for a non-spike time index, 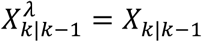. Whether there is a spike on time index *k* or not, 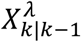 is defined by a mixture of Gaussians. The last step in the filter update is multiplying 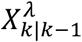 by exp(−Δ_*k*_ Λ(*X*_*k*_)).

Multiplication of 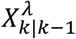 by exp(−Δ_*k*_ Λ(*X*_*k*_)) does not follow a Gaussian mixture model structure; however, we can use Gaussian approximation method to represent the posterior using a Gaussian mixture model. The other possible solution is to first approximate exp(−Δ_*k*_ Λ(*X*_*k*_)) using a Gaussian mixture model; and then the filter update follows a similar procedure already described in deriving 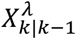 on a spike time. Note that exp(−Δ_*k*_ Λ(*X*_*k*_)) is the same on all filtering time-steps, and it can be approximated by a MoGs once for the entire processing time. This methodology becomes problematic if we use many mixture components to approximate exp(−Δ_*k*_ Λ(*X*_*k*_)); this is because, the number of mixture components become significantly large within a few filtering time-steps, so we choose to approximate the Gaussian approximation method to build the posterior [43]. Using this method, the number of Gaussian mixture components does not grow – we refer to 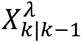, and it is a reasonably accurate approximation specifically when Λ(*X*_*k*_) doesn’t have many local maxima and sharp curvatures. This happens when there are many cells firing over the maze space a rat explores. Let’s assume 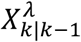 is defined by

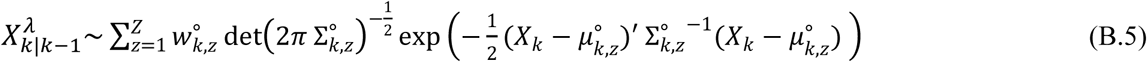

where *Z* might be either V_*k*-1_ U or V_*k*-1_ and 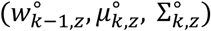 are the mixture components parameters. To update the weight, mean, and covariance of each mixture component, we Taylor expand - Δ_*k*_ Λ(*X*_*k*_), the logarithm of exp(−Δ_*k*_ Λ(*X*_*k*_)), about a different point for each mixture component. Specifically, for the *z*^*th*^ mixture component, we expand the - Δ_*k*_ Λ(*X*_*k*_) about the one-step prediction mean 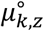 for that component using a second-order Taylor expansion. Finally, we complete the square to generate a new GMM. We use the following updates for the posterior mean, covariance, and mixture weight of each mixture component

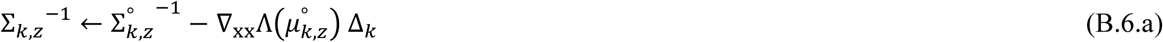

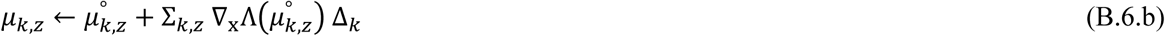

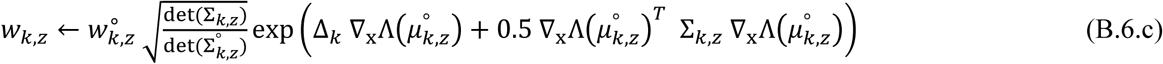

where, ∇_x_ and ∇_xx_ are the gradient and Hessian operators [44]. Note that there is a closed-form solution for the gradient and Hessian operators for a normal density function and GMM. The gradient and Hessian for a normal density function with (*μ*, ∑) at point *X* is defined [42] by

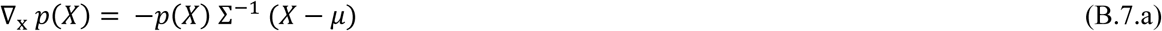

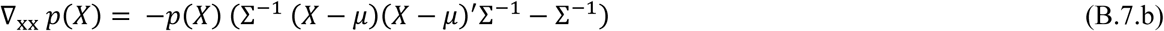

where, *p*(*X*) is the density function of a multivariate normal with (*μ*, ∑) parameters. Note that in (B.6.a), there is a possibility of an updated covariance matrix to become non-positive definite. In this case, we ignore updating this covariance estimate and thus its mean and weight update. In other words, we only update those mixture components, for which their updated covariance matrix is positive definite (PSD).

Here, we described every processing step needed in calculating the filter update rule at each filtering time-step. This provides a closed-form solution for a marked point process filter when the joint mark intensity function is defined by a MoGs. The posterior distribution generated here will be the input to our dropping and merging algorithms which are designed to optimally control the growth of mixture components over consecutive processing time.

## Appendix C Performance Analysis of Dropping and Margining Algorithms using a KL divergence measure

KL divergence [26] is widely used in machine learning and particularly distribution approximation [45]. Here, we analyzed the performance result of our proposed algorithm using a KL distance in 1-D decoding, already introduced in the application section. The problem setup and modeling parameters are the same, and we only use the *KL*(*P*||*Q*) in place of *B*(*P*||*Q*) to assess similarity between *P*(*X*) and *Q*(*X*) distributions.

**Figure C.1** shows the decoded trajectory using KL divergence method with *α*_*d*_ = 0.15 and *α*_*d*_ = 0.12. The graph shows that the decoder fails to follow the rat trajectory. Also, the posterior distribution shows a larger growth over space compared to the exact method. We examined different values of *α*_*d*_ and *α*_*m*_, and we observed similar characteristics there as well. We ran the same performance analysis described in the application section, and the performance results are shown in **Figure C.2** and **table C.1** respectively. The result shows that we reach a better computational efficiency as the average number of mixtures becomes lower using the KL distance. For instance, for *α*_*d*_ = 0.15 and *α*_*d*_ = 0.12, the average computational saving is about 10% better than the drop-merge method using *B*(*P*||*Q*). However, we observed a larger RMSE error and a significant reduction in 95% HPD measure.

As **Figure C.1** shows, the decoder fails to follow the rat movement trajectory. However, we can argue when the decoder follows the trajectory, we can get a better 95% HPD coverage area performance; however, this comes with a pay off in RMSE error. This implies that we expect to have a larger RMSE using KL divergence, even when it properly traces the rat movement trajectory.

**Figure C.1.**
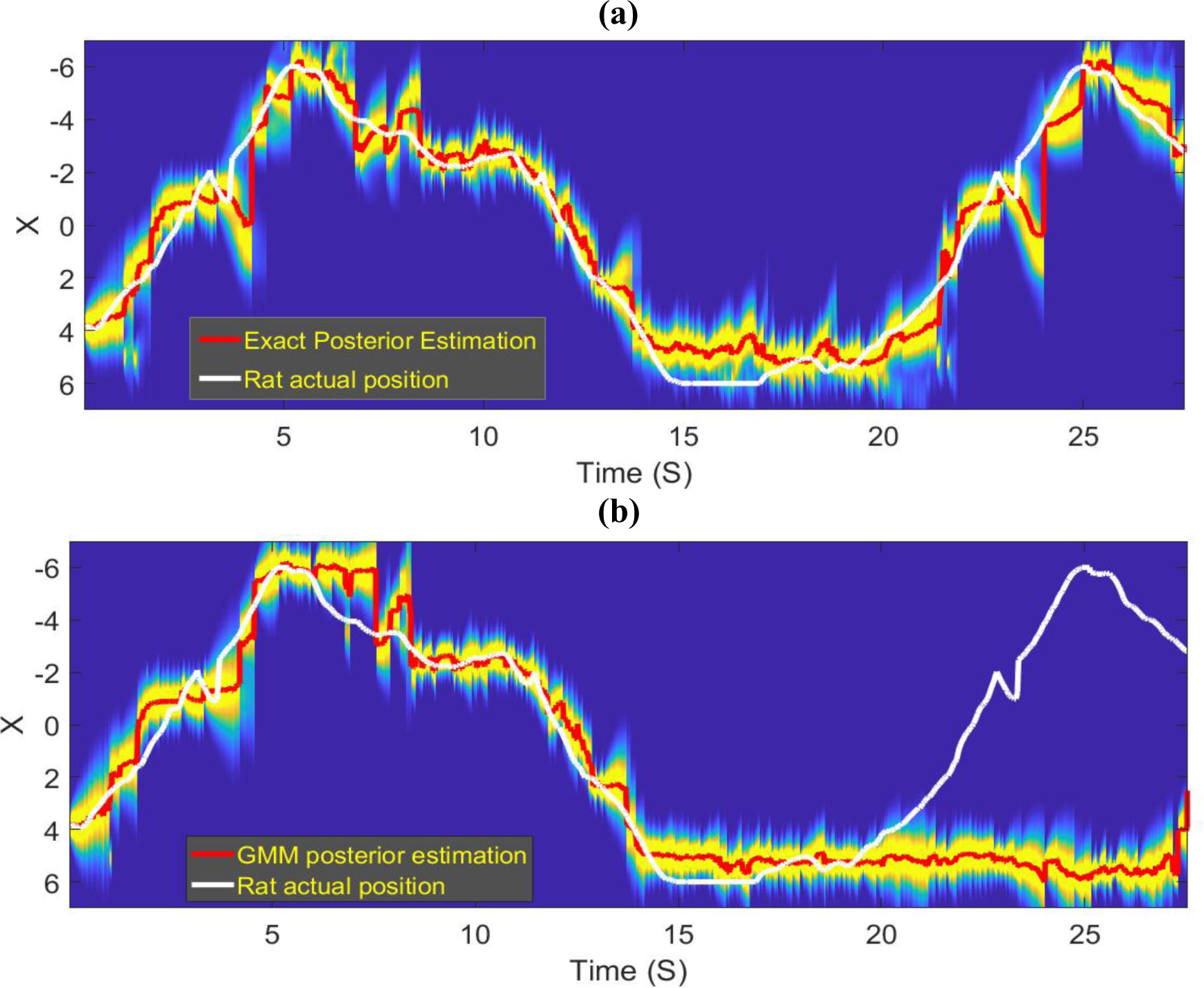
Decoding result using KL divergence **a.** Decoding result using the exact solution (this is the same as **figure 2(a)**) **b.** Decoding result using KL divergence method with *α*_*d*_ = 0.15 and *α*_*m*_ =012. Decoding result using drop-merge method with KL divergence measure follows the rat movement trajectory for the first half of the data, but it fails to properly trace the movement trajectory for the second half of the data.

**Table C1.** Performance result using the exact and proposed solution in 1-D decoding problem.

| Method | Setting | RMSE<br>(cm) | 95%<br>HPD | Processing<br>Time<br>(msec) | Avg.<br>Number of<br>Mixtures | Average Number<br>of Mixtures on<br>Spike Time | Maximum<br>Number of<br>Mixtures |
| --- | --- | --- | --- | --- | --- | --- | --- |
| Exact Solution | $dx=0.025$ | 0.80 | 80.5 | 24.4 | NA | NA | Na |
| GMM-based<br>method | $\alpha_d = 0.15$<br>$\alpha_m = 0.12$ | 4.2 | 55.7 | 1.0 | 1.08 | 2.48 | 9 |

**Figure C.2.**
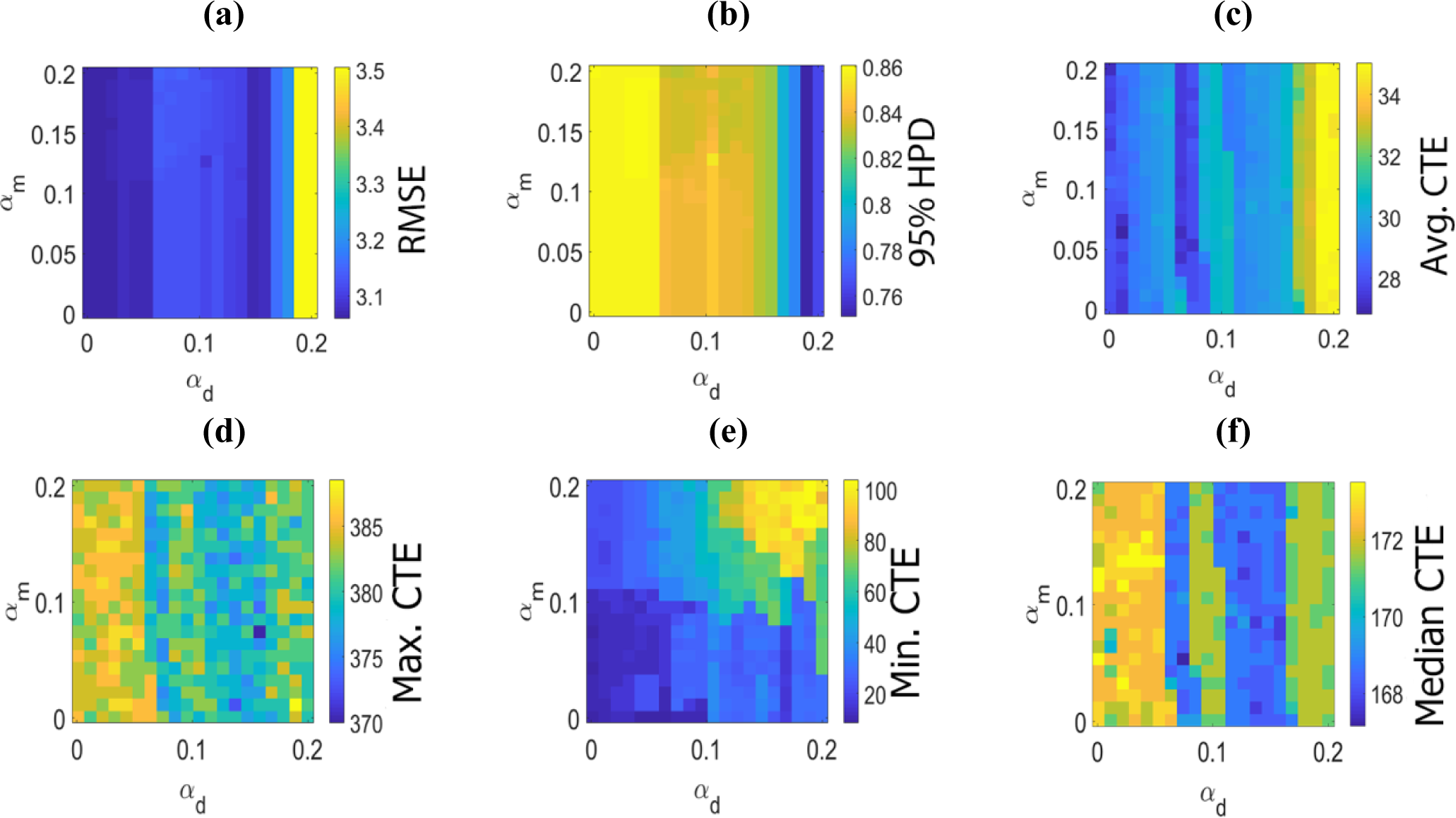
Performance and computational time efficiency of KL divergence method for 1-D decoding task using different *α*_*d*_ and*α*_*m*_ parameters. **b.** 95% HPD coverage performance map. **c**. Average computational time efficiency (CTE) using KL drop-merge method. **d.** Maximum computational time efficiency using KL drop-merge method. **e.** Minimum computational time efficiency using KL drop-merge method. **f.** Median of computational time efficiency using KL drop-merge method.

## Appendix D Further Analysis on Computation Efficiency and Performance of the Exact and Drop-merge Solution

We argued that the processing time of the drop-merge algorithm changes as the number of mixture components in GMM being passed to these algorithms change. Note that the number of mixture components in these GMMs are different given a spike event happens or not. Thus, we study how the drop-merge algorithm processing time change as a function of the number of mixture components in GMMs at spike and non-spike time events.

**Figure D.1(a)** and **figure D.1(b)** show the average and the maximum number of mixture components being passed to the drop-merge algorithm per each time-step. For *α*_*m*_ = 0.12 and *α*_*d*_ = 0.15, this number are 1.25 and 12 respectively. **Figure D.1(e)** shows the average computational time saving on the spike times; for *α*_*m*_ = 0.12 and *α*_*d*_ = 0.15, the processing time of the drop-merge algorithm on spike time is about the same for the exact solution. **Figure D.1(f)** shows the average computational time saving on non-spike times; for *α*_*m*_ = 0.12 and *α*_*d*_ = 0.15, the computational time saving is about 2500. This result suggests that computational saving is not merely emerging from utilizing the merging and dropping algorithms on each filtering time-step; however, there is a significant computational saving on the non-spike time where the merge and drop algorithms deal with a relatively low number of mixture components in GMM -note that merging or dropping are not needed when there is one mixture component for filter solution in the previous time point. Non-spike times are about 95% of the whole processing time, and this is being reflected in the overall computational saving. Note that we gain this computational saving because we build a parametric distribution model – here, a GMM – for the posterior distribution of the rat position on each filtering time step, and this leads to a super-fast algorithm – even, analytical solution – when the number of mixture components in GMM becomes small.

**Figure D.2** provides further information on the processing time for specific numbers of mixture components in GMMs being passed to the drop-merge algorithm. Note that the number of mixture components in the conditional intensity of the cell ensembles plays a role in the number of mixture components being generated in these GMMs, which are not studied here. For the MoG model used in the 1-D decoding task, the number of mixture components is 35. This means on the spike times, the number of mixture components in the GMM being passed to the drop-merge algorithms are 35 times higher than non-spike times.

**Figure D.3** shows the histogram of processing time over the whole dataset. These graphs are consistent with the results shown in **figure D.1(e)** and **figure D.1(f).** The processing time for 80% of time points are less than 1 millisecond using the drop-merge algorithm.

**Figure D.1.**
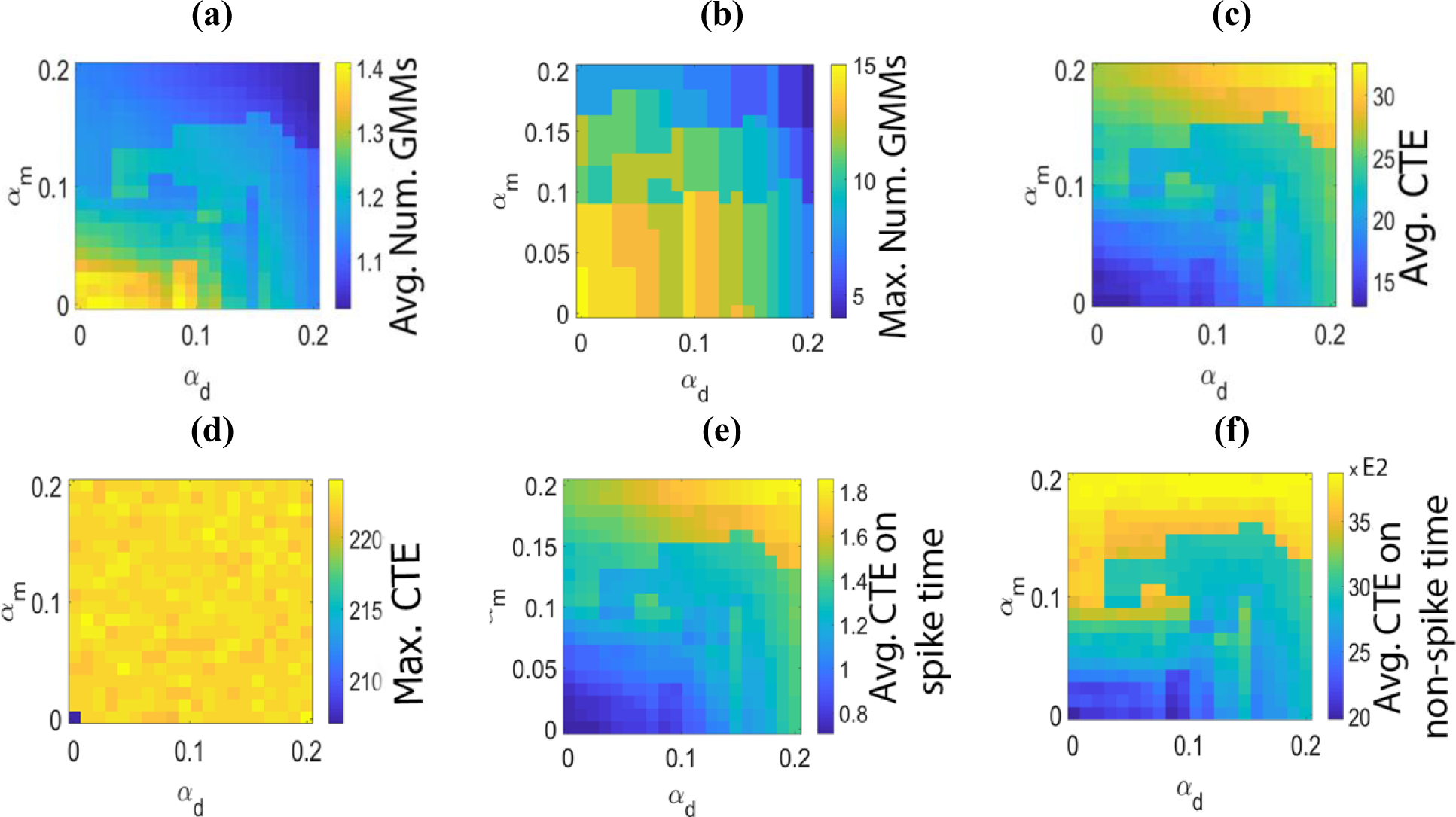
Computational time analysis and its relationship with the number of mixture models in drop-merge method for 1-D decoding task. **a.** Average number of GMM components for different values *α*_*d*_ and *α*_*m*_ created for the filter solution at each processing time step. **b.** Maximum number of GMM components for different values *α*_*d*_ and *α*_*m*_ created for the filter solution at each processing time step. **c**. Average computational time efficiency using the drop-merge method. Here, the average processing time in the exact method is divided by the average processing time per time step using the drop-merge algorithm. So, a value of 25 implies that the drop-merge method run 25 times faster than the exact solution. **d.** Maximum computational time efficiency in the drop-merge method. For instance, a value of 215 implies that for the corresponding parameter setting there is at least one time step where the computation saving is 215 times faster than average processing time of the exact solution. **e.** Average computational time efficiency using the drop-merge method on spike times. Here a value of 1.5 implies that the drop-merge method run 1.5 times faster than the exact solution on spike times. **f.** Average computational time efficiency using the drop-merge method on non-spike time point. For example, a value of 3000 implies that the drop-merge method run 3000 times faster than the exact solution on non-spike times.

**Figure D.2.**
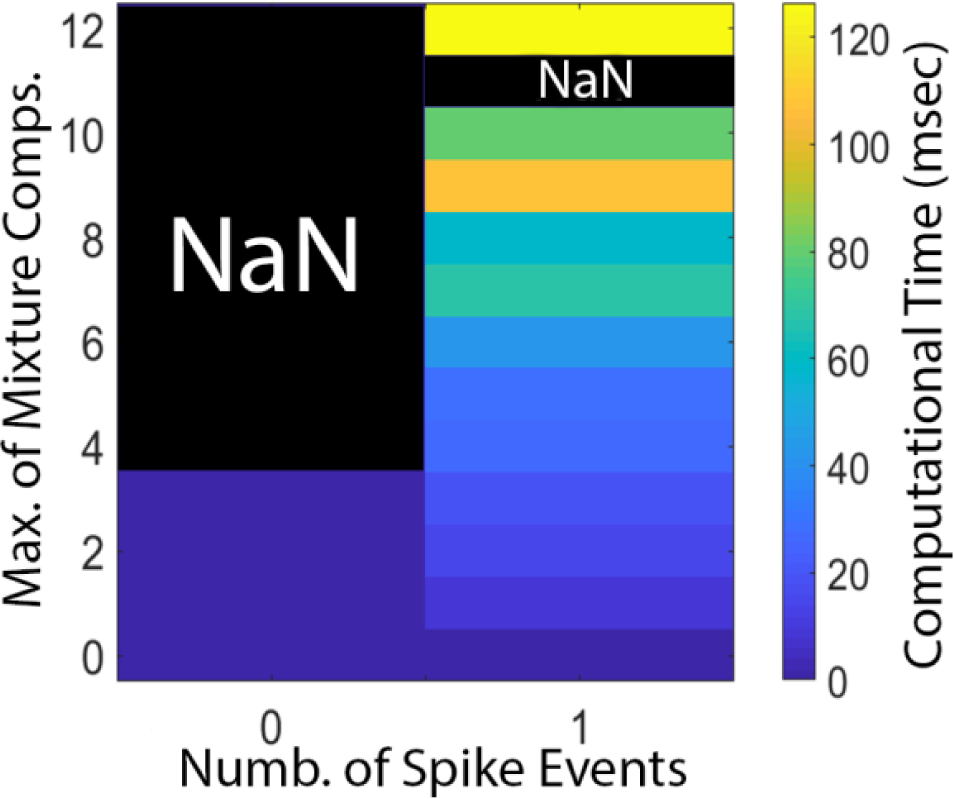
1-D decoding task computational time in (ms) for different number of mixture models per spike and non-spike time. The merge and drop parameters are *α*_*d*_ = 0.15 and *α*_*m*_ = 0.12. Note the average processing time for the exact solution is 22 msec. Note that Y-axis - number of mixture components - represents number of mixture components on the previous time filter solution; merge and dropping algorithms process 35 times of this number.

**Figure D.3.**
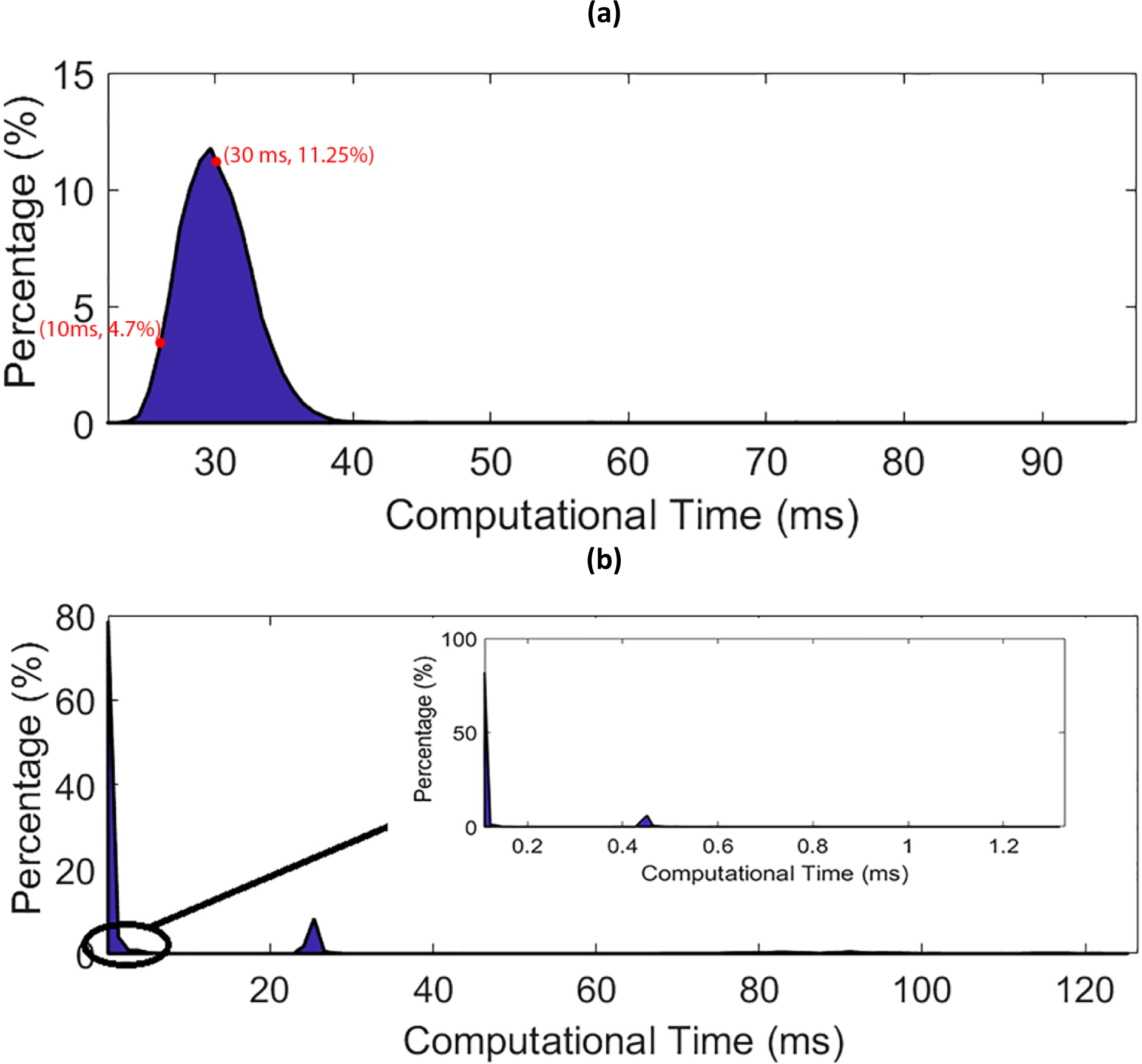
Histogram of 1-D task computational time for the exact filter solution and drop-merge method. **a.** Histogram of computational time for the exact filter solution. Its mean is about 25 msec, and as it shown most of timesteps have computation time near this value. **b.** The histogram of computational time for drop-merge method with α_d_ = 0.15 and α_m_ = 0.12. We provide a closer look to computational time for a processing time below 1.2 msec. The result shows, most of computational times for 1-D task in the drop-merge algorithm are below 1.0 msec. This means the algorithm is real-time in most timesteps. Also, we can see there are some timsteps with computational time near 23.0 msec, about 8.0% of timesteps, and less than 0.1% of timesteps with computational time more than 60 msec.

The result in 2-D decoding showed a significant boost in computational time efficiency compared to 1-D decoding. Here, we repeated the same sort of analysis run for 1-D decoding for 2-D decoding. **Figure D.4(a)** and **figure D.4(b)** suggest the number of mixture components in the GMMs being passed to the drop-merge algorithm is relatively lower than 1-D decoding. This means that the overall computational time efficiency will be even larger compared to 1-D decoding. **Figure D.4(e)** and **figure D.4(f)** support this assumption; **figure D.4(e)** shows that we even get the computational time efficiency at spike times – a property which was absent in 1-D decoding. **Figure D.4(f)** shows that this computational time efficiency is huge at non-spike time points, and thus, we reach a computationally efficient algorithm for 2-D decoding. **Figure D.5** and **Figure D.6** will further support this claim and provide a further information about the merging-dropping algorithm characteristics. Note that the histogram of the computational time in the exact solution is in second time scale.

**Figure D.4.**
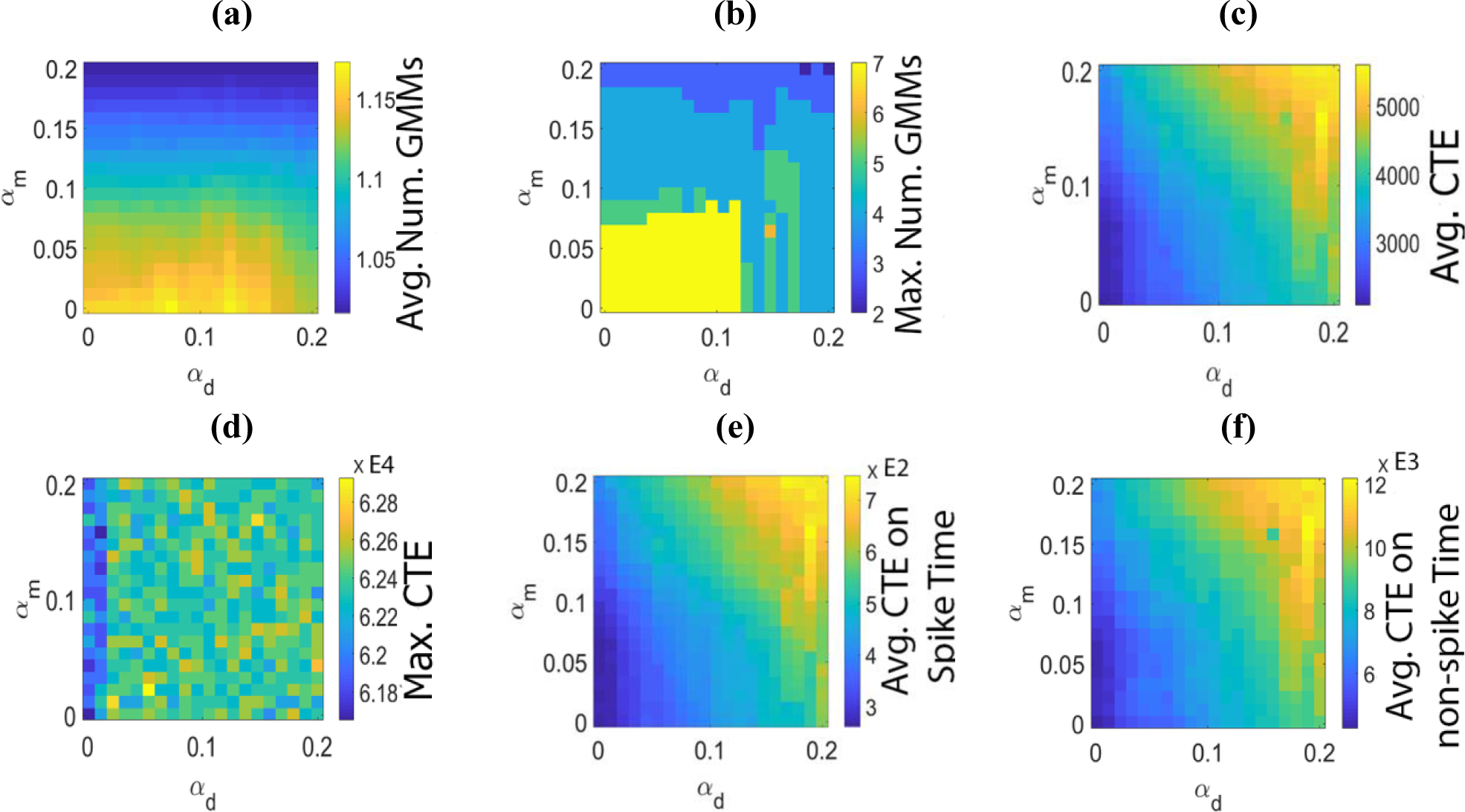
Computational time efficiency analysis and its relationship with the number of mixture models in drop-merge method for 2-D decoding task. **a.** Average number of GMM components for different values *α*_*d*_ and *α*_*m*_. **b.** Maximum number of GMM components for different values *α*_*d*_ and *α*_*m*_. **c**. Average computational time efficiency using the drop-merge method. Here, the average processing time in the exact method is divided by the average processing time per time step using the drop-merge algorithm. So, a value of 4000 implies that the drop-merge method run 4000 times faster than the exact solution. **d.** Maximum computational time efficiency in the drop-merge method. For instance, value of 62000 implies that for corresponding sets of parameters, there are time steps that runs about 62000 times faster than the exact solution. **e.** Average computational time efficiency using the drop-merge method on spike times. Here a value of 500 implies that the drop-merge method run 500 times faster than the exact solution on spike times. **f.** Average computational time efficiency using the drop-merge method on non-spike time point. For example, a value of 8000 implies that the drop-merge method run 8000 times faster than the exact solution on non-spike times.

**Figure D.5.**
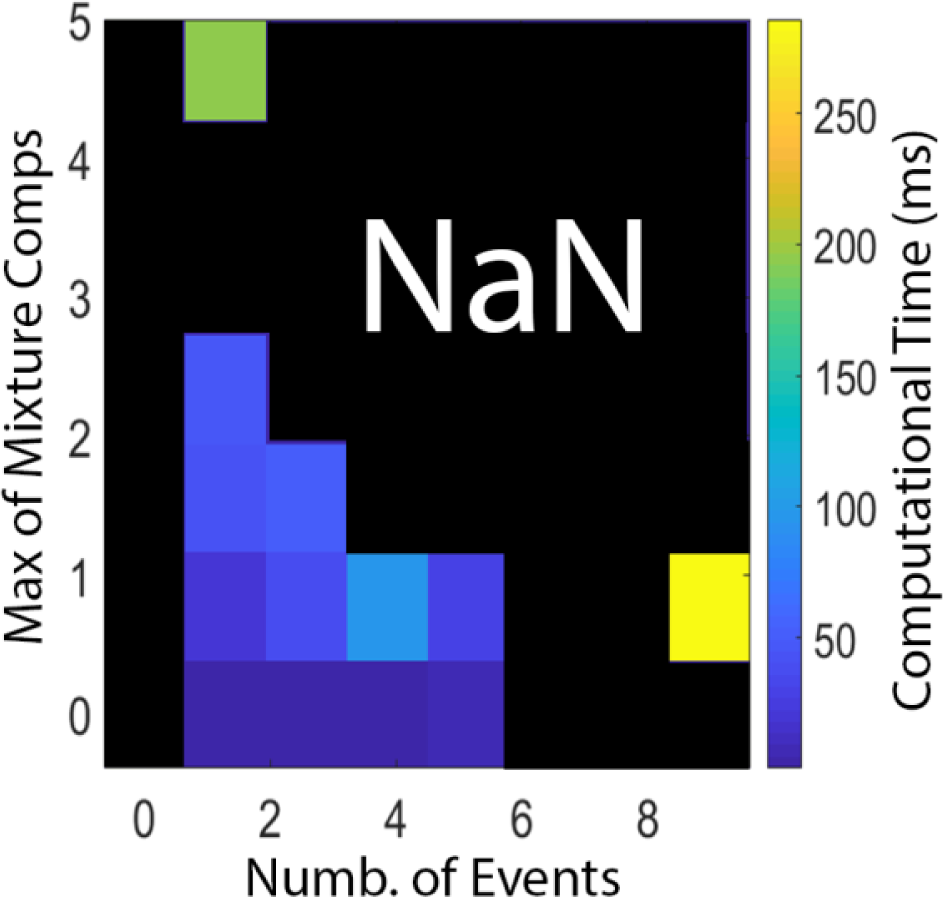
2-D decoding task Computational time result (ms) with α_d_ = 0.05 and α_m_ = 0.1 parameters setting for different combinations of mixture models and number of spikes in drop-merge method. Columns of the figure describe maximum number of mixture models, and rows describe number of spikes. Each element of the figure is the average computational time for related combination of number of spikes and maximum number of mixture models.

**Figure D.6.**
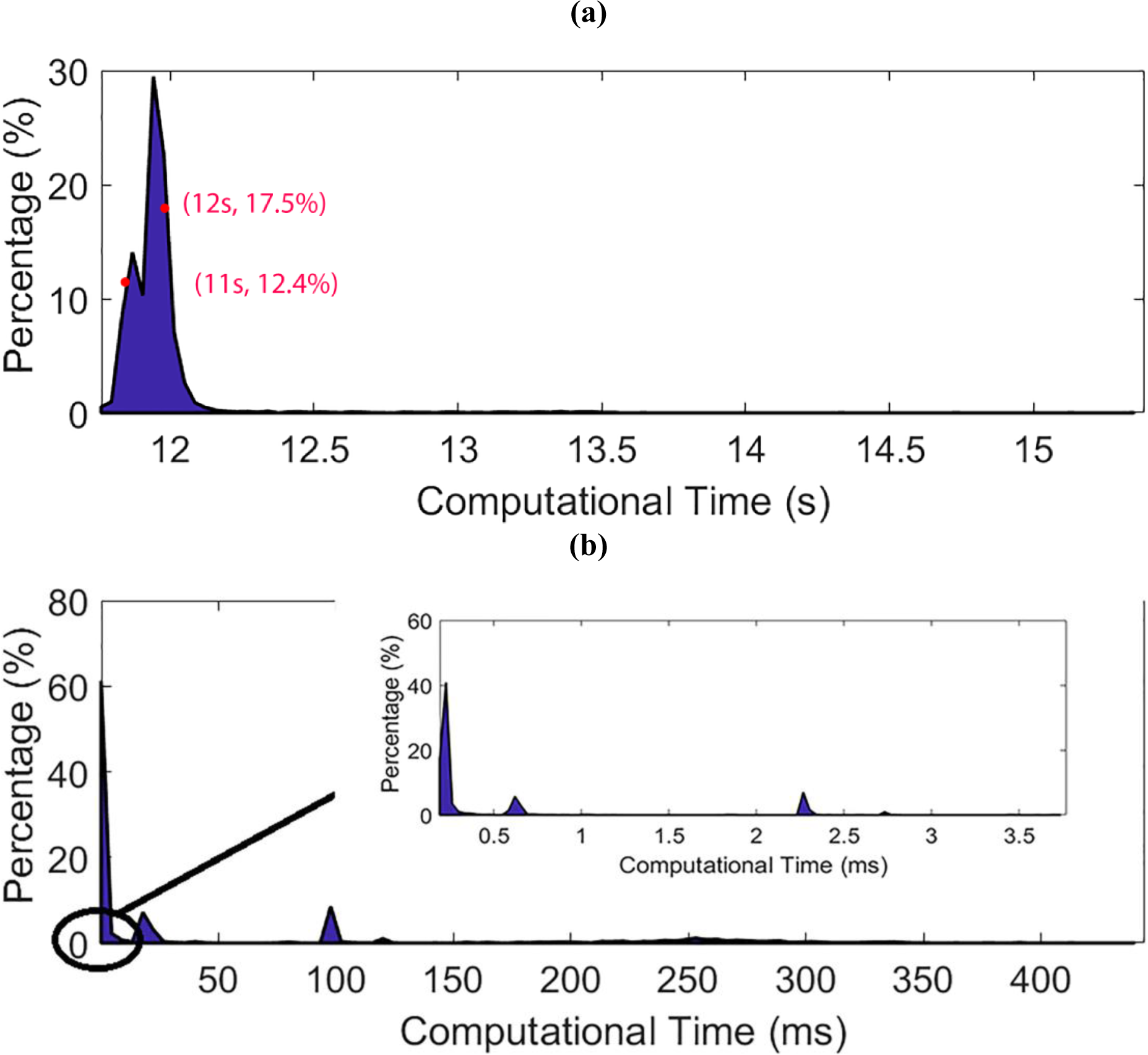
Histogram of 2-D task Computational time for exact filter solution and drop-merge method. a. histogram for exact filter solution. Its mean is about 11.5 sec, and as it shown most of timesteps have computation time near this value. b. histogram for drop-merge method with α_d_ = 0.05 and α_m_ = 0.1 parameters setting. We provide a closer look to computational time for a processing time below 3.5 msec. The result shows, most of computational times for 2-D task in the drop-merge algorithm are below 1.0 msec. This means the algorithm is real-time in most timesteps. Also, we can see there are some timsteps with computational time near 100.0 msec, about 9.1% of timesteps, and less than 0.1% of timesteps with computational time more than 300 msec.

